# Macrophage differentiation is marked by increased abundance of the mRNA 3’ end processing machinery, altered poly(A) site usage, and sensitivity to the level of CstF64

**DOI:** 10.1101/2022.08.07.503082

**Authors:** Srimoyee Mukherjee, Joel H. Graber, Claire L. Moore

## Abstract

Regulation of mRNA polyadenylation is important for response to external signals and differentiation in several cell types, and results in mRNA isoforms that vary in the amount of coding sequence or 3’ UTR regulatory elements. However, its role in differentiation of monocytes to macrophages has not been investigated. Macrophages are key effectors of the innate immune system that help control infection and promote tissue-repair. However, overactivity of macrophages contributes to pathogenesis of many diseases. In this study, we show that macrophage differentiation is characterized by shortening and lengthening of mRNAs in relevant cellular pathways. The cleavage/polyadenylation (C/P) proteins increase during differentiation, suggesting a possible mechanism for the observed changes in poly(A) site usage. This was surprising since higher C/P protein levels correlate with higher proliferation rates in other systems, but monocytes stop dividing after induction of differentiation. Depletion of CstF64, a C/P protein and known regulator of polyadenylation efficiency, delayed macrophage marker expression, cell cycle exit, attachment, and acquisition of structural complexity, and impeded shortening of mRNAs with functions relevant to macrophage biology. Conversely, CstF64 overexpression increased use of promoter-proximal poly(A) sites and caused the appearance of differentiated phenotypes in the absence of induction. Our findings indicate that regulation of polyadenylation plays an important role in macrophage differentiation.

## 1 Introduction

Macrophages are a critical component of the innate immune system that have a protective and beneficial role through the phagocytosis and killing of microbes, removal of dying cells and cellular debris, production of immune-regulatory molecules, and presentation of antigens to T-cells (1). Macrophages are derived from precursor cells called monocytes, which, in turn, are derived from myeloid progenitors. Monocytes undergo phenotypical and functional differentiation when they are recruited to tissues, where they differentiate into macrophages with altered morphological features and then engage in inflammatory or tissue remodeling processes (2). Robust macrophage function is critical for tissue repair and combating infection. However, inappropriate or overly robust activity contributes to the pathogenesis of diseases such as atherosclerosis, arthritis, asthma, lung fibrosis, type 2 diabetes, insulin resistance, inflammatory bowel disease, cancer and most recently, COVID-19 and the formation of intra-abdominal scar tissue (3–9).

Abnormal proliferation of macrophage progenitor cells in lieu of differentiation not only impairs normal macrophage responsibilities but can also lead to blood cancers such as acute myeloid leukemias (AML). A promising approach to treat AML is differentiation therapy, in which malignant cells are encouraged to differentiate into more mature forms using pharmacological agents. This approach has been remarkably successful for a small subset of AML cases involving a PML-retinoic acid receptor fusion (10, 11), and is a valuable alternative approach to killing cancer cells through cytotoxicity. To improve treatment of leukemias by forcing differentiation as well as to modulate macrophage function to help abate diseases influenced by macrophages, we must understand the critical molecular pathways that govern macrophage differentiation.

In many cell types, differentiation requires not only changes in transcription, but also pervasive changes in mRNA splicing and 3’ end cleavage and polyadenylation (C/P). In the C/P step, pre-mRNA is cleaved at the poly(A) site and a poly(A) tail is added to the new end (12). Changes in polyadenylation play an increasingly appreciated role in regulation of gene expression by affecting protein output at many levels. For example, the efficiency at which a particular mRNA precursor is polyadenylated will affect the amount of mRNA available for translation. Through alternative polyadenylation (APA), genes with multiple poly(A) sites can produce distinct mRNA isoforms with different lengths of 3’ untranslated regions (3’ UTRs), which in turn affects nuclear export, translation, stability, association with stress granules, and localization in the cytoplasm (13–16). Alternate sequences in the 3’ UTR can even facilitate co-translational protein complex formation (17) or act as a sink to bind miRNAs that down-regulate tumor suppressor mRNAs (18). Polyadenylation that occurs in an intron or less frequently an exon upstream of the 3’ most exon affects the coding region, leading to expression of different protein isoforms (19, 20).

Many examples of APA have been documented in the immune system. For example, stimulation of human monocytes with lipopolysaccharide (LPS) and interferon-γ or activation of B cells and T cells leads to proliferation and a switch to upstream poly(A) sites (21, 22). A specific example of the importance of APA in immune cell differentiation is the switch to the upstream poly(A) site of the transcription factor NFATC1, which leads to NFATC1 accumulation and maturation of naïve T cells into effector T cells (23). Preferential utilization of the upstream poly(A) site of the immunoglobulin heavy chain gene in activated B cells is needed to switch to producing the secreted form of the protein instead of the membrane-attached form and is mediated by the CstF64 C/P protein (24–26). Increased levels of CstF64 and changes in APA have also been observed after treatment of RAW 264.7 mouse monocyte/macrophage-like cells with LPS (27), which activates these cells to produce an inflammatory response (28).

While the above studies have illuminated key aspects of mRNA 3’ end processing in immune cell regulation, the potential role of C/P proteins in differentiation of monocytes to macrophages, which results in a dramatic change of cellular characteristics including attachment to surfaces, increased size with extended filopodia, and cell cycle arrest, has not been explored. We hypothesized that the changes in cell phenotypes that occur during macrophage differentiation will be accompanied by changes in choice of poly(A) site, and that perturbation of the C/P machinery will affect the course of this differentiation. In the current study, we show that this is indeed the case. There are significant changes in APA when the monocytic U937 cell line was differentiated to form macrophages in vitro, as well as an increase in the components of the C/P machinery. Genetic manipulation of the C/P protein CstF64 led to altered APA and differentiation properties of monocytes. Overall, our findings indicate an important role for regulation of polyadenylation in macrophage differentiation.

## 2 Results

### 2.1 Establishment of the monocyte-macrophage differentiation systems

To study alternative polyadenylation in monocyte to macrophage differentiation, we used the well-established monocytic cell lines U937 and THP-1 (29) as models. These cells can be differentiated to macrophages by treatment with phorbol 12-myristate 13-acetate (PMA) (30) and are commonly used to study macrophage differentiation and function. While the two cell lines have different origins, with THP-1 being derived from a patient with Acute Myeloid Leukemia and U937 from a patient with histiocytic lymphoma (29), they allow investigations using cells with a homogeneous genetic background. After 6 hours of treatment with 30 nM of PMA, U937 cells are in a transition phase in which they stop dividing and loosely adhere to the dish, with little change in morphology (Fig. 1A). By 18-24 hours, almost all cells showed macrophage-like phenotypes including increased cell size, strong adherence to the dish, and appearance of filopodia. Monocyte-macrophage differentiation is associated with decreased expression of monocyte surface markers and increased expression of macrophage markers (31). The classic macrophage surface markers such as CD16 (32), CD68 (33), HLA-DRA (34) and CD38 (35) were considerably increased over the 24-hour time course and the monocyte marker CD14 (36) had reduced expression in differentiated U937 cells (Fig. 1B). As observed previously (37, 38), these cells also displayed increased attachment and no longer proliferated after PMA treatment (Fig. 1C and 1D). THP-1 cells behaved similarly to the U937 cells in terms of morphological changes, marker expression, attachment, and proliferation (Supplementary Fig. S1A-D).

**Figure 1.**
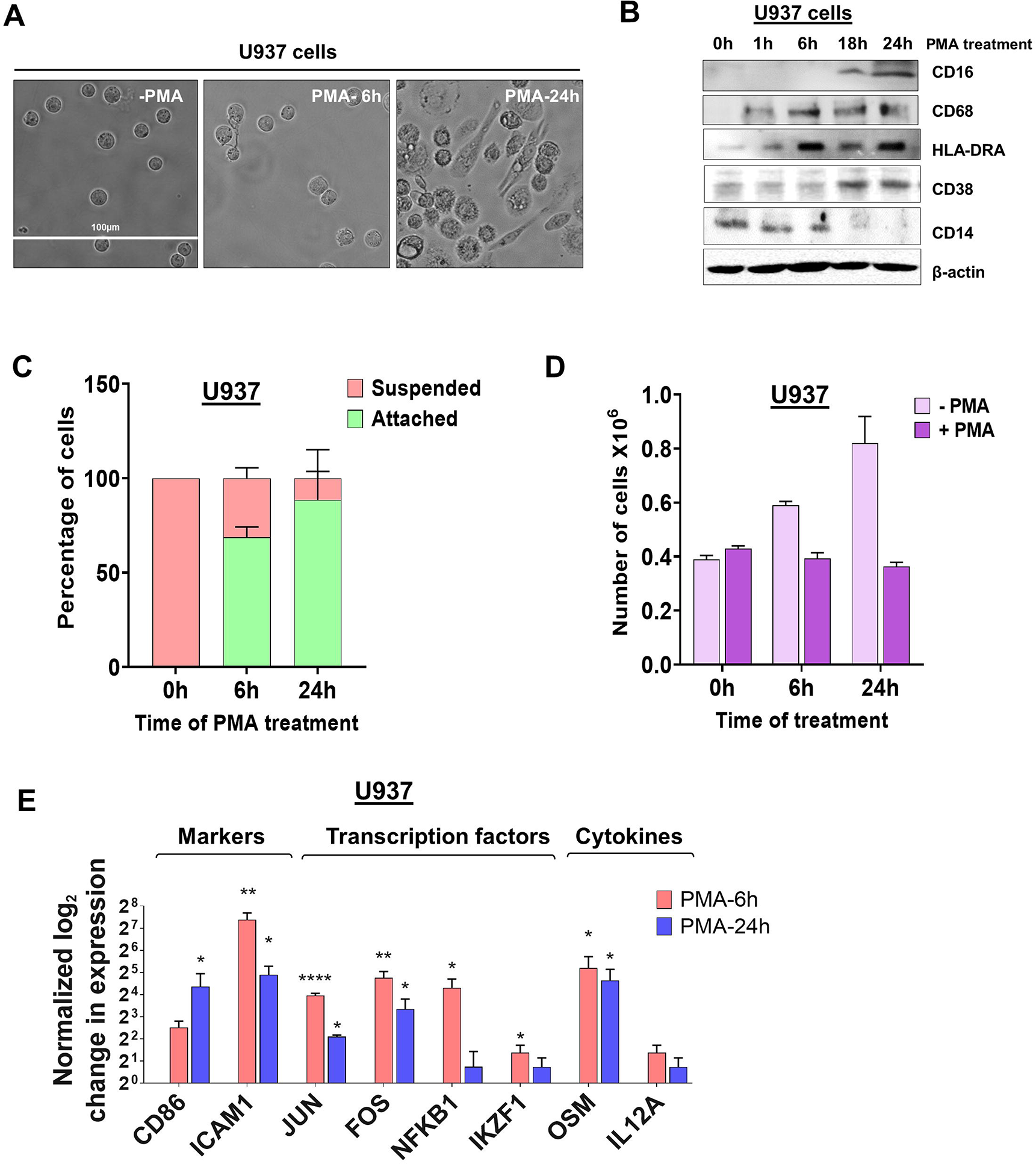
Morphological changes and altered expression of cell-surface markers and mRNAs during monocyte differentiation. (A) The effect of differentiation on cellular morphology. U937 cells exposed to PMA (30 nM) for 6 and 24 hours and observed by phase contrast microscopy (40X). (B) Western blots of total cell lysates from U937 cells differentiated with PMA (30 nM) for 0, 1, 6, 18, and 24 hours. Blots were probed with antibodies against CD16, CD68, HLA-DRA, CD38 and CD14 with β-actin as the loading control. (C) Attachment assay of U937 cells. Cells were treated with PMA for 0, 6, and 24 hours and live cells visualized and counted by the Trypan blue exclusion assay. The graph presents the percentage of cells that are suspended or attached. (D) Proliferation assay of U937 cells. The number of cells before and after differentiation depicts the absolute cell number in millions, compared to normal cell proliferation in the absence of PMA. The figure represents mean ± SE from three independent experiments. (E) Real-time quantitative PCR (RT-qPCR)–based analysis of the expression of genes important for macrophage differentiation and function *(CD86, ICAM1, JUN, FOS, NFKB1, IKZF1, OSM* and *IL12A)*. The RT-qPCR analysis used primers designed in the first exon to represent the total transcript level and the figure shows log_2_-fold changes in expression of these genes at 6 hours and 24 hours post PMA treatment of U937 cells, where values are normalized to ACTB mRNA and the 0 hour (no PMA) time point. The figure represents mean ± SE from three independent experiments. P value <0.05 was considered significant, where * = P ≤ 0.05; ** = P ≤ 0.01; *** = P ≤ 0.001; **** = P ≤ 0.0001.

To further validate that differentiation was successful in our systems, we measured the mRNA levels of genes important for macrophage differentiation and function at two critical time points in the transition from monocytes to macrophages - 6 hours after induction when U937 and THP-1 cells show some signs of differentiation and 24 hours, when differentiation is complete. Genes encoding proteins such as the surface marker CD86, the adhesion molecule ICAM1, transcription factors including JUN, FOS, NFKB1, and IKZF1, cytokines such as Oncostatin-M (OSM) and IL12A, all increased in expression at 6 hours and some also at 24 hours after treating U937 and THP-1 cells with PMA (Fig. 1E and Supplementary Fig. S1E).

### 2.2 Integrative analysis of poly(A) site mapping data identifies expression changes and 3′ UTR shortening and lengthening events during macrophage differentiation

To determine the effects of PMA treatment on global poly(A) site usage and gene expression in U937 cells, we used Poly(A)-Click sequencing, which returns sequences adjacent to the poly(A) site and has been used in recent studies to assess APA and expression changes (39–43). The differential expression for a particular gene can be determined by collapsing all mapped poly(A) sites within each gene into one count. As might be expected for monocytes undergoing differentiation, genes with increased expression at 6h are strongly enriched in pathways involved in the immune system and macrophage function (Supplementary Fig. S2A). Pathways associated with DNA repair were predominant in the decreased expression set. At 24 hours and similar to what was seen at 6 hours, the categories with significant increases in expression were heavily skewed towards the immune system and pathways related to macrophage function (Supplementary Fig. S2B). The increased expression of macrophage-related genes further confirms that our differentiation protocol is effective. The down-regulated set at 24 hours also showed marked enrichment of genes involved in cell cycle and chromosome organization. Decreased expression of these genes is consistent with the exit from the cell cycle upon PMA treatment.

We also asked if there were changes in poly(A) site usage during differentiation. The degree of APA usage was calculated by changes in the Percentage of Distal poly(A) site Usage Index (delta-PDUI), which identifies lengthening (positive index) or shortening (negative index) of 3′ UTRs having statistical significant difference between differentiated and control samples. For U937 cells differentiated for 6 hours, sequencing identified polyadenylated mRNAs from a total of 12,208 genes of which 331 genes had mRNAs with significant changes in length in the terminal 3’ UTR of the 3’ most exon, designated as terminal 3’ UTR. Of these, 139 exhibited an increase in shorter isoforms, and 113 showed an increase in longer isoforms, while 79 showed a complex pattern involving multiple poly(A) sites, where use of some went up and others went down (Fig. 2A). At 24 hours, signals from 12,145 genes were obtained, and mRNAs from a greater number of genes (864) had changes in poly(A) site usage compared to 6 hours. Of these, 376 exhibited shortening, 305 exhibited lengthening and 183 showed a complex pattern (Fig. 2A). At both time points, the number of significantly shortened 3’ UTR APA events was slightly higher compared to the significantly lengthened APA events. Some genes have poly(A) sites located upstream of the gene’s last exon. Use of these intronic sites creates a new terminal exon and changes the amount of coding sequence in the mRNA. We also observed changes in intronic poly(A) site use, but at a lower frequency than changes in the terminal 3’ UTR. At 6 hours, 9 genes showed increased use of these poly(A) sites and one showed decreased use, and at 24 hours, 45 genes showed an increase and 30 showed a decrease (Supplementary Fig. S5A and B). Detailed outputs for all the APA events are supplied in Supplementary Tables S3-S5.

**Figure 2.**
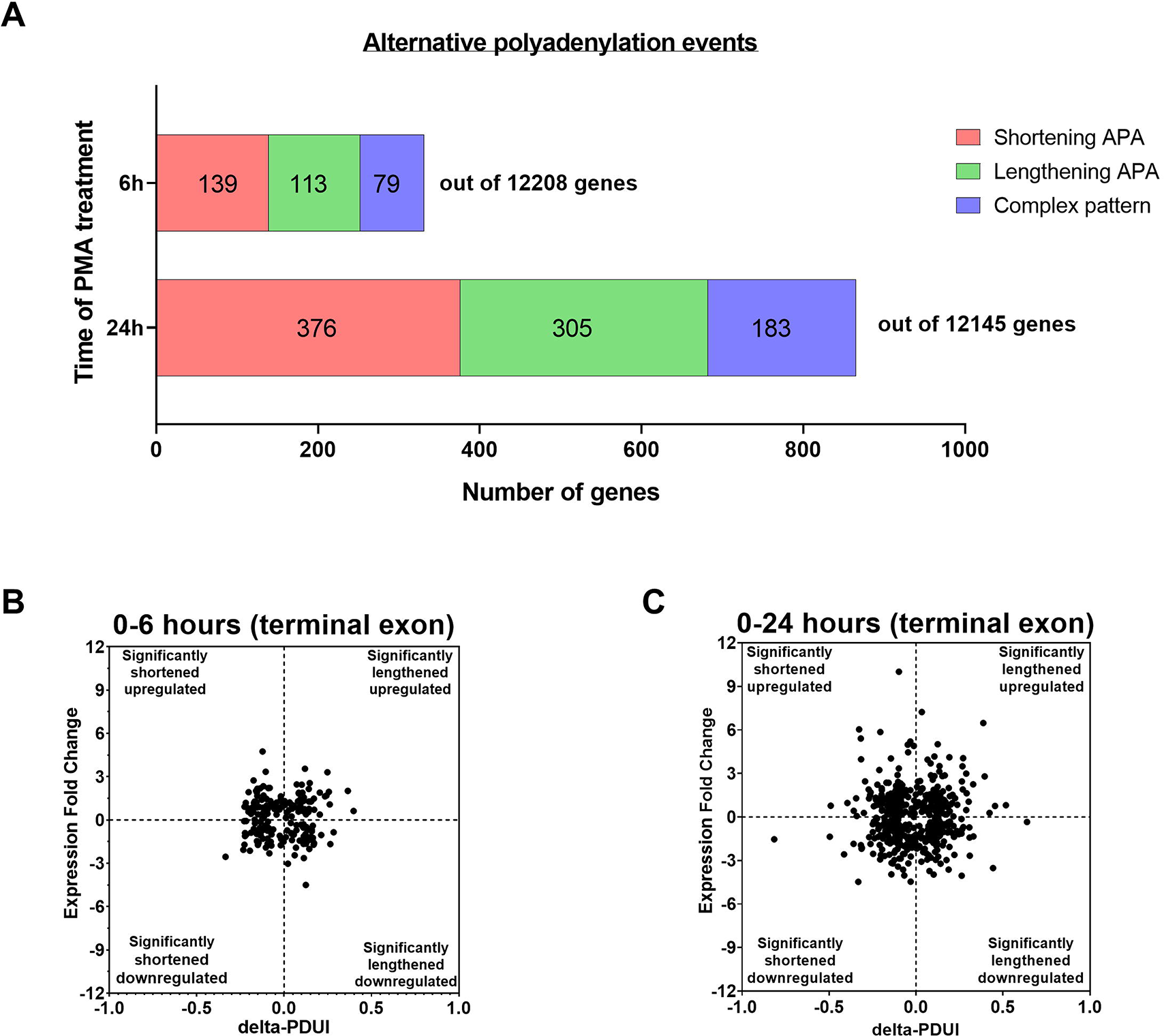
Evidence of APA in U937 cells after 6 and 24 hours of PMA treatment. (A) Bar graph representing the numbers of genes with APA changes in the terminal 3’ UTR after 6h and 24h of PMA treatment with respect to control undifferentiated U937 cells. Red, green and blue represents shortening, lengthening and complex (where some went up and others went down) APA patterns in the dataset. The criteria for a significant APA change is that at least one PAC in an exon must change greater than 1.5 fold with p_Adj_ <0.1, resulting in a difference in fractional usage of that PAC of at least 10%. (B & C) Log_2_-fold change in gene expression is plotted against the change in Percentage of Distal poly(A) site Usage Index (delta-PDUI) for 3′-UTR-altered genes in 6h PMA-treated U937 cells with respect to control undifferentiated ones (B) or in 24h PMA-treated U937 samples (C). Plots for fold change vs poly(A) site usage (delta PDUI) where genes on the left half of each graph represent 3′-UTR-shortened hits, whereas those on the right half represent 3′-UTR lengthened hits.

Shortening of transcripts in the terminal 3’ UTR has sometimes, but not always been associated with increased mRNA expression, with the converse for mRNA lengthening (22, 44-49). To assess how gene expression correlated with changes in terminal 3’ UTR length during macrophage differentiation, the 3’ UTR shortened and lengthened gene sets were categorized into those with up-regulated or down-regulated expression and delta-PDUI versus fold change in expression was plotted for the 6 and 24 hour time points (Fig. 2B and Fig. 2C). A comparable number of genes in the lengthened or shortened groups at both time points exhibited increases or decreases in mRNA expression, and many genes with significant changes in isoform length showed no or only small changes in expression, indicating a poor correlation between 3’ UTR length and steady-state mRNA level.

Previous work has shown that genes with significant changes in their 3’ UTR isoform expression ratios are cell type- and pathway-specific (48). To determine the biological pathways impacted by APA during macrophage differentiation, we performed Gene Set Enrichment Analysis (GSEA) (50) on genes which exhibited significant shortening or lengthening of transcripts. Analysis of the 0 vs 6h shortened set showed an enrichment of categories related to immune system or signal pathways related to macrophage function, as well as metabolism and cellular transport (Fig. 3A). Intracellular transport is important for both cytokine secretion and phagocytosis, and modulation of metabolism is recognized as a central player in macrophage activation (51). In contrast, the lengthened set was primarily related to the immune system.

**Figure 3.**
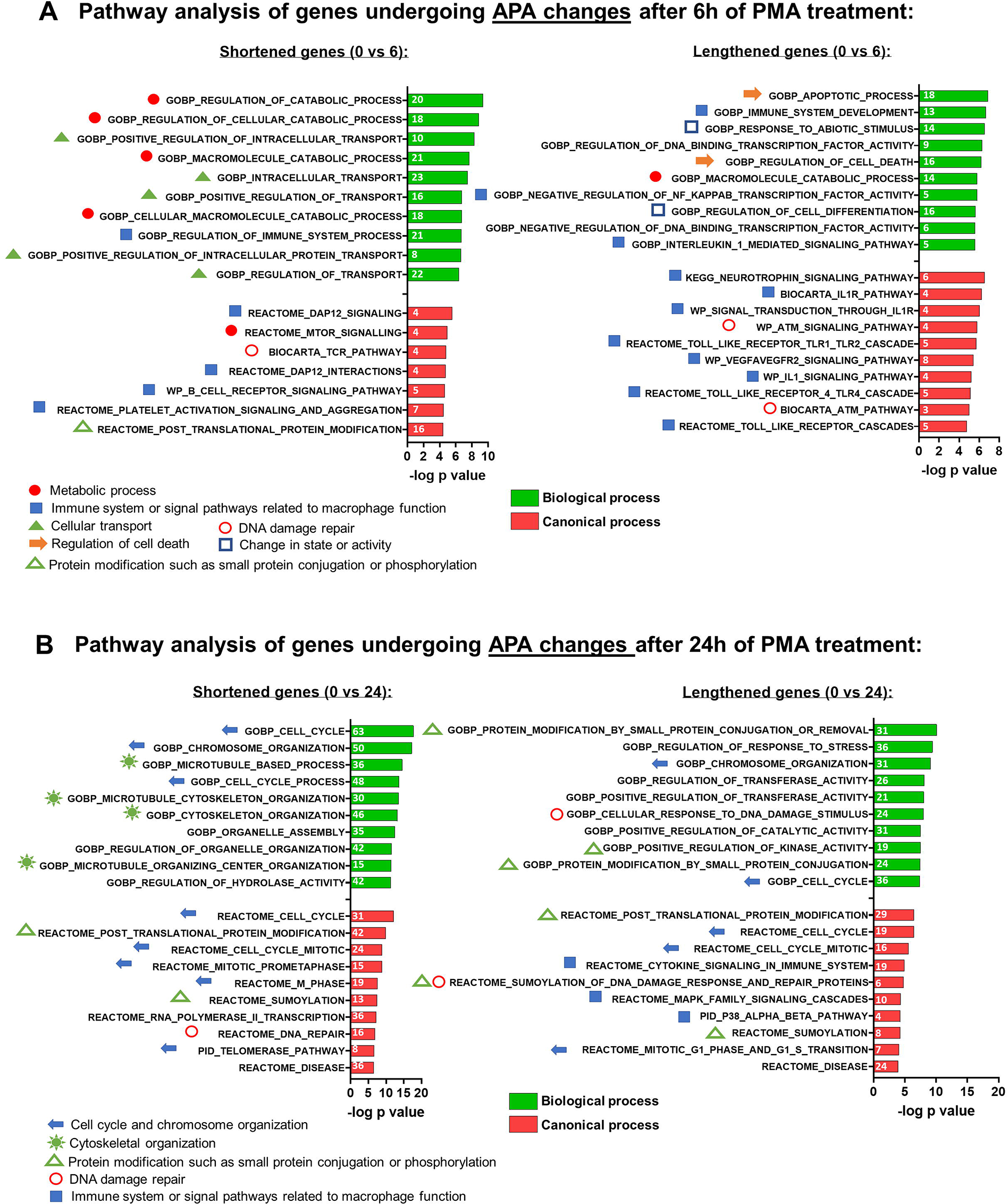
Gene set enrichment analysis (GSEA) of APA events in U937 cells after PMA treatment. Functional annotation clustering of mRNAs whose APA is regulated and after 6h (A) and 24h (B) of differentiation. The 10 most significant GO-process enriched gene groups for biological process (green) or canonical process (red) are ranked based on negative log_10_ (P-values) for shortened (left panel) and lengthened (right panel) transcripts. Pathways classified according to specific cellular processes are grouped together as indicated by the individual symbols at the bottom of the graphs.

The affected pathways for the 0 vs 24h APA changes were quite different from those affected at 6 hours. Unlike what was observed for 6h, APA changes at 24h that involved shortening affected pathways not specific to the immune system but nevertheless relevant to macrophage differentiation such as cytoskeletal organization, cell cycle, and chromosome organization, while the lengthened set was more mixed and included categories related to post-translational modification, cell cycle and signaling pathways involved in macrophage-mediated inflammatory responses (Fig. 3B).

Examples of genome browser plots of the mapped poly(A) sites for genes which showed mRNA shortening or lengthening in U937 cells are shown in Supplementary Fig. S3A for the 6 hour time point (CIAPIN1 and EIF1) and in Fig. 4 for the 24 hour time point (shortened: SERPINB1 and PTCH1; lengthened: PSAT1 and MRPL44). One way to confirm the presence of isoforms differing in their 3’ ends and shifts in use of these isoforms is 3’ rapid amplification of cDNA ends (RACE) assays (52–55). Thus, 3’ RACE assays were used to detect the different mRNA isoforms, and these confirmed the APA changes depicted by the genome browser plots from the poly(A)-sequencing data of U937 cells (Fig. 5 and Supplementary Fig. S3B). SERPINB1 has 3 major poly (A) sites (PAC-1, 3 and 5 as shown in Fig. 4A), but the 3’ RACE was not able to detect the longest isoform, perhaps because of the strong bias towards amplifying the short isoform limited the sensitivity for longer products. Nevertheless, in accordance with the genome browser plot, we did observe a reduction in use of the middle p(A) site (marked as D in Fig. 4A). The differentiation of the other monocytic cell line, THP-1 also produced shortening or lengthening of the same genes after 24 hours of PMA treatment (Supplementary Fig. S4A and B).

**Figure 4.**
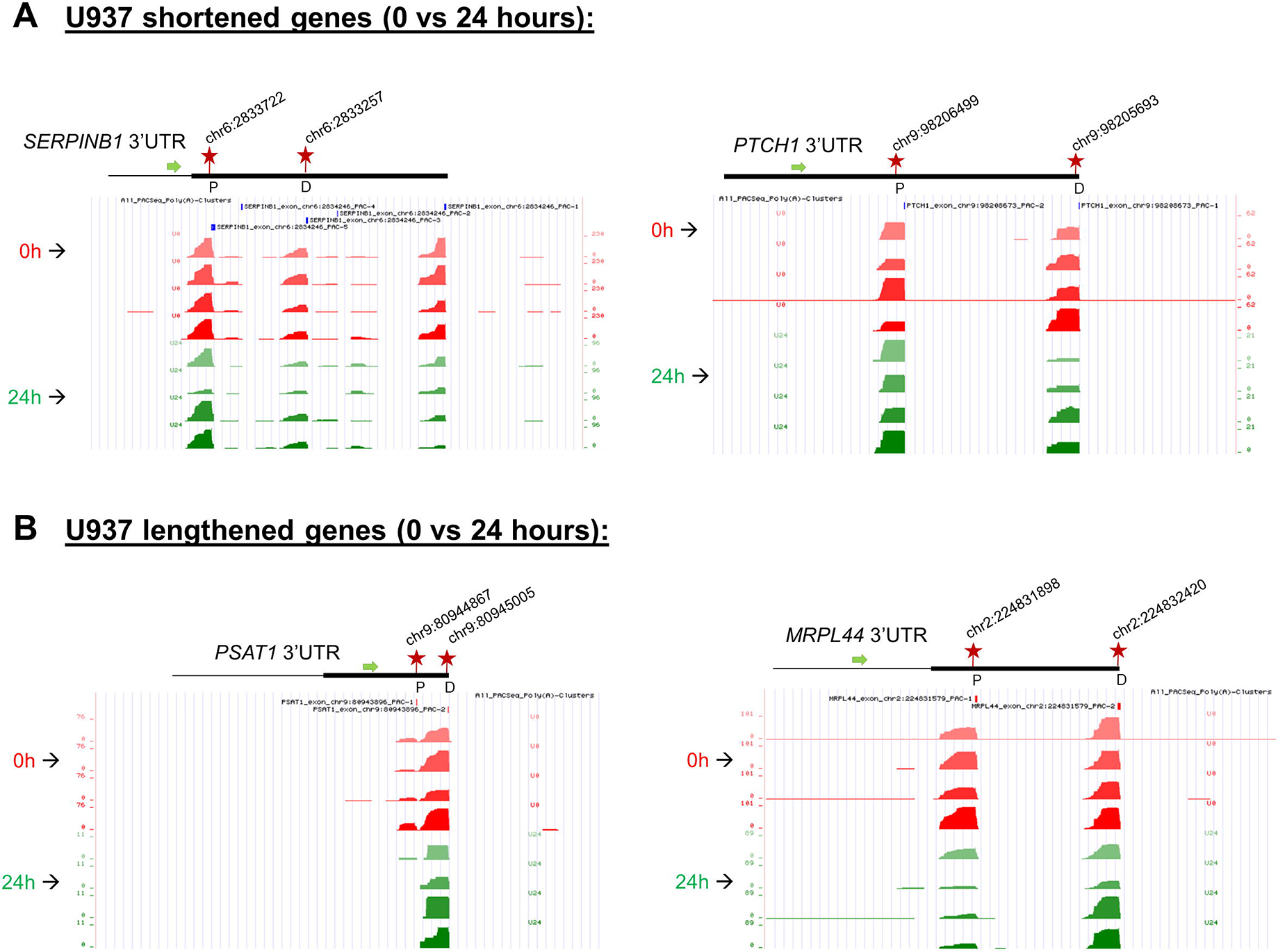
Changes in poly(A) site use in U937 cells after 24 hours of PMA treatment. UCSC genome browser plots of RNA sequencing tracks highlighting the 3′-UTR profile differences for shortened genes (A) or lengthened genes (B) after 24 hours of PMA treatment with respect to control (0h). The colors of the tracks represent 0h (red) and 24h (green). Proximal (P) and distal (D) poly(A) sites are indicated with red stars. The green arrow defines the direction of the coding strand, and tag counts are indicated on the y axis. Additionally, positions and chromosome co-ordinates of the annotated PACs are indicated at the top of each plot.

**Figure 5.**
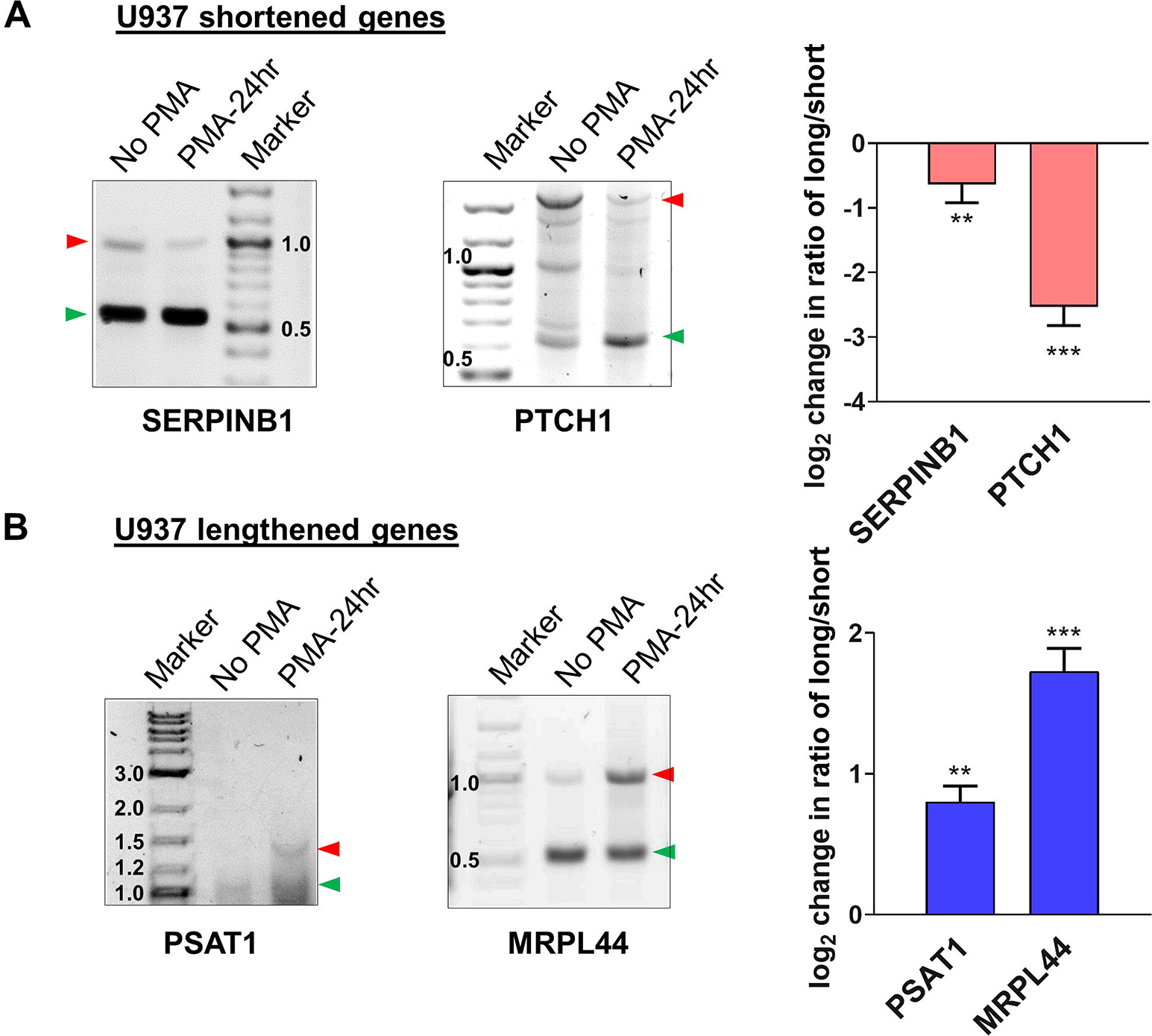
Validation of APA events in U937 by 3’ RACE. Representative 1% agarose gel images for the 3′ RACE RT-PCR from U937 cells (untreated and 24h PMA-treated) for (A) shortened genes SERPINB1, PTCH1 and (B) lengthened genes PSAT1 and MRPL44 using gene specific primers at the last exon-exon junction and a common reverse anchor primer. To confirm the direction of the shift in poly(A) site use, normalized intensities of long and short bands for each PCR were quantified and a long/short ratio determined and normalized to the No PMA control. The corresponding graphs represent mean ± SE from at least two independent experiments. P value <0.05 was considered significant, where * = P ≤ 0.05; ** = P ≤ 0.01; *** = P ≤ 0.001.

### 2.3 Levels of cleavage/polyadenylation (C/P) complex proteins are upregulated at different times during monocyte-to-macrophage differentiation

The analysis of the poly(A)-Click seq data described above shows that the APA occurring during monocyte-macrophage differentiation affected relevant functional categories of genes, and that similar numbers of genes showed shortening or lengthening in the 3’ UTR, with a slight trend towards shortening. We next wanted to explore possible mechanisms for eliciting this APA and examined the C/P complex proteins, as numerous studies have shown that changing the levels of these proteins can affect poly(A) site choice (56, 57). The core C/P complex is comprised of six factors: CPSF, CstF, CFI_m_, CFII_m_, symplekin and poly(A) polymerase (PAP) (Fig. 6B) [reviewed in (58)]. The C/P proteins, with the exception of PAP, increased in level during differentiation of U937 cells, but with different kinetics and not necessarily in the same manner for the individual subunits of each factor (Fig. 6A). For example, CstF50, CstF64 and CstF77 increased early at 1 hour after PMA treatment, before differentiated phenotypes are observable. CPSF100, CPSF73 and CFI_m_59 increased at 6 hours when cells begin to attach, whereas CFI_m_68, FIP1, CPSF160, Clp1, Pcf11, symplekin and CstF64τ, a conserved paralog of CstF64 (59) in mammals, increased later (18-24 hours), when the cells are mostly differentiated and attached. Some subunits, such as WDR33 and CstF77, decreased when differentiation was completed. The C/P proteins increased in similar ways during differentiation of the THP-1 monocytic cell line, with comparable kinetics (Supplementary Fig. S6A). Overall, our findings show that C/P proteins increase during macrophage differentiation but that some may be limiting in concentration at different points in the differentiation process.

**Figure 6.**
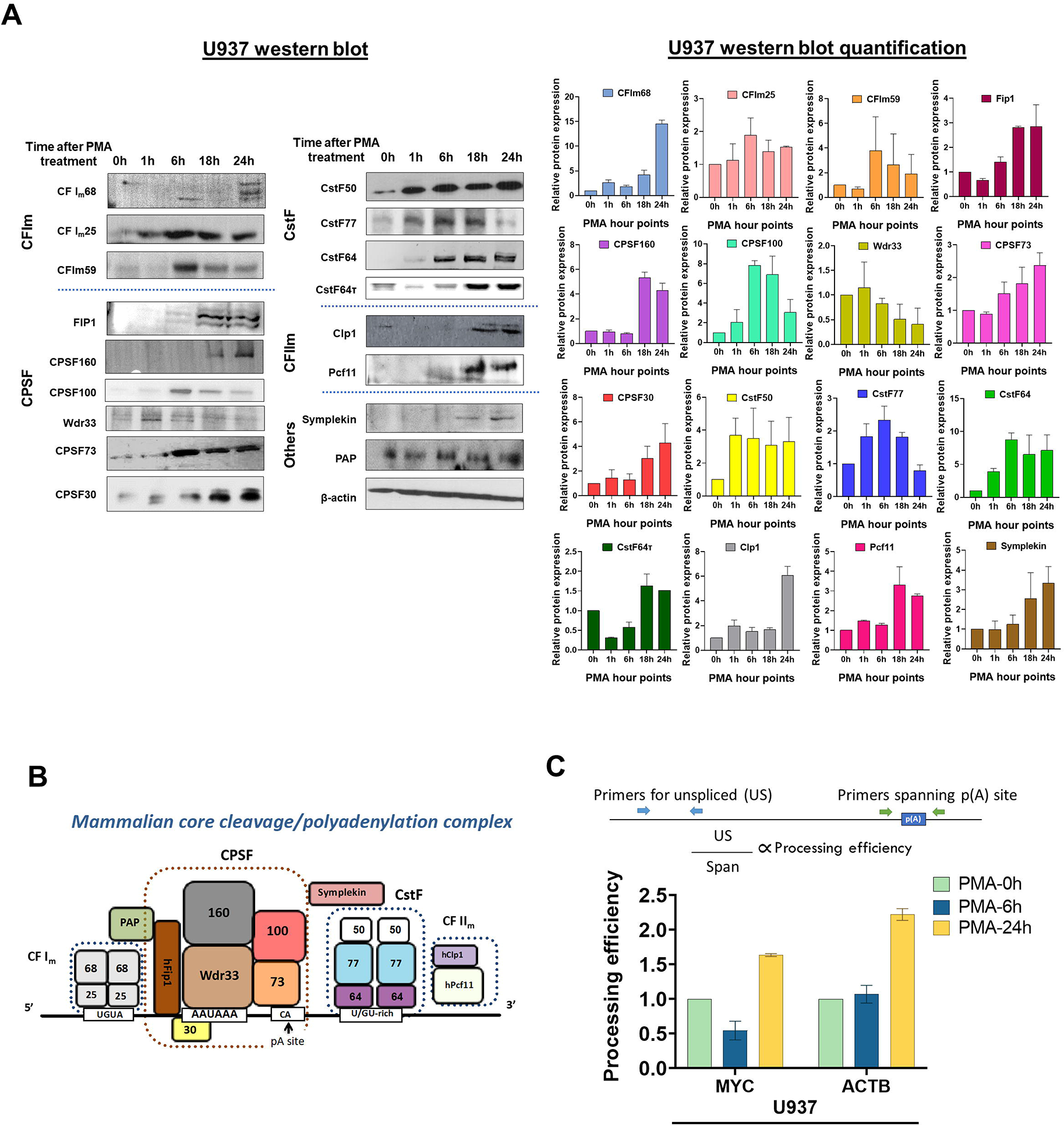
Effect of macrophage differentiation on the expression of C/P proteins in U937 cells. (A) Whole cell lysates from U937 cells treated with PMA for 0, 1, 6, 18 and 24 hours were separated by 10% SDS-PAGE and western blotting was performed for the indicated subunits of the C/P complex, with β-actin as the loading control. Each western blot (left panel) was performed in three biological replicates of the differentiation process and the quantified data is shown in the right panel. (B) Schematic of the mammalian core C/P complex. (C) Processing efficiency of MYC and ACTB transcripts for U937 cells. The bar graph shows the ratio of unspliced (US) RNA transcripts (detected by RT-qPCR with a primer pair upstream of the poly(A) site) to unprocessed RNA (detected by RT-qPCR with a primer pair that spans the poly(A) site) as shown in the schematic. The cDNA preparation was done using random hexamers.

### 2.4 Differentiation of monocytic cells increases processing efficiency at single poly(A) sites

To test whether PMA-mediated differentiation alters the efficiency of 3’-end processing *in vivo*, we determined the level of uncleaved transcripts that will contain sequence upstream and downstream of a poly(A) site. We used previously described primers to monitor cleavage of ACTB or MYC transcripts in vivo by RT-qPCR of total RNA reverse transcribed with random hexamers (60, 61). ACTB has a single poly(A) site and MYC has a major poly(A) site used 93% of the time (62). One primer pair detects RNA containing sequence that spans the poly(A) sites (Span) of the ACTB or MYC genes. Because we expect uncleaved RNA to be nuclear, we compared this value to that obtained using a primer pair which detects unspliced transcripts (US), which would also be nuclear. The processing efficiency at each gene’s poly(A) site can be quantified by the ratio of “US” PCR product to that of “Span” product. If processing efficiency increases, fewer “Span” products are expected relative to the “US” control Fig. 6C (61). The processing efficiency at each gene’s poly(A) site can be quantified by the ratio of US PCR product to that of Span product. Compared to undifferentiated cells, more mRNA from both genes was processed at 24 hours, the time when most of the U937 or THP-1 cells have differentiated (Fig. 6C and Supplementary Fig. S6B). Respective “minus RT” controls gave insignificant signals (Supplementary Fig. S7A) indicating the effect to be entirely coming from the transcripts and not DNA contamination. This data indicates that consistent with the increase in C/P proteins, differentiation positively affected processing efficiency at these ACTB and MYC poly(A) sites.

### 2.5 Genetic manipulation of CstF64 alters monocyte-macrophage differentiation and switches poly(A) site choice

As described above, monocyte differentiation led to the increased expression of C/P proteins and altered APA of mRNAs from groups of genes known to be important for macrophage differentiation and function. We thus wanted to test whether altering the level of one of these proteins could impact the APA and macrophage differentiation. We focused on CstF64, because it has been shown to influence APA during several types of transitions in cell state, in particular some of which involve the immune system (21-23, 27, 63, 64). Based on these previous studies, we hypothesized that while CstF64 is likely not to be the only C/P protein affecting APA in macrophage differentiation, alteration of its levels would affect both APA and the progression of differentiation.

To test our hypothesis, we generated stable cell lines in which the expression of CstF64 in U937 cells could be knocked down (KD) by doxycycline-inducible expression of shRNA against CstF64 mRNAs. Depletion of CstF64 was done by doxycycline treatment for 3 days and then PMA was added for one day before harvesting the cells. Assessment of the extent of protein KD by two different shRNA sequences against CstF64 showed considerable decrease in expression, indicating efficient KD, while levels of the homolog CstF64τ remained unaffected (Fig. 7A). The macrophage surface markers CD16, CD68, HLA-DRA and CD38 were substantially reduced compared to the control shRNA when either CstF64 shRNA was used for KD. On the other hand, the levels of monocyte marker CD14 increased on CstF64 KD (Fig. 7A).

**Figure 7.**
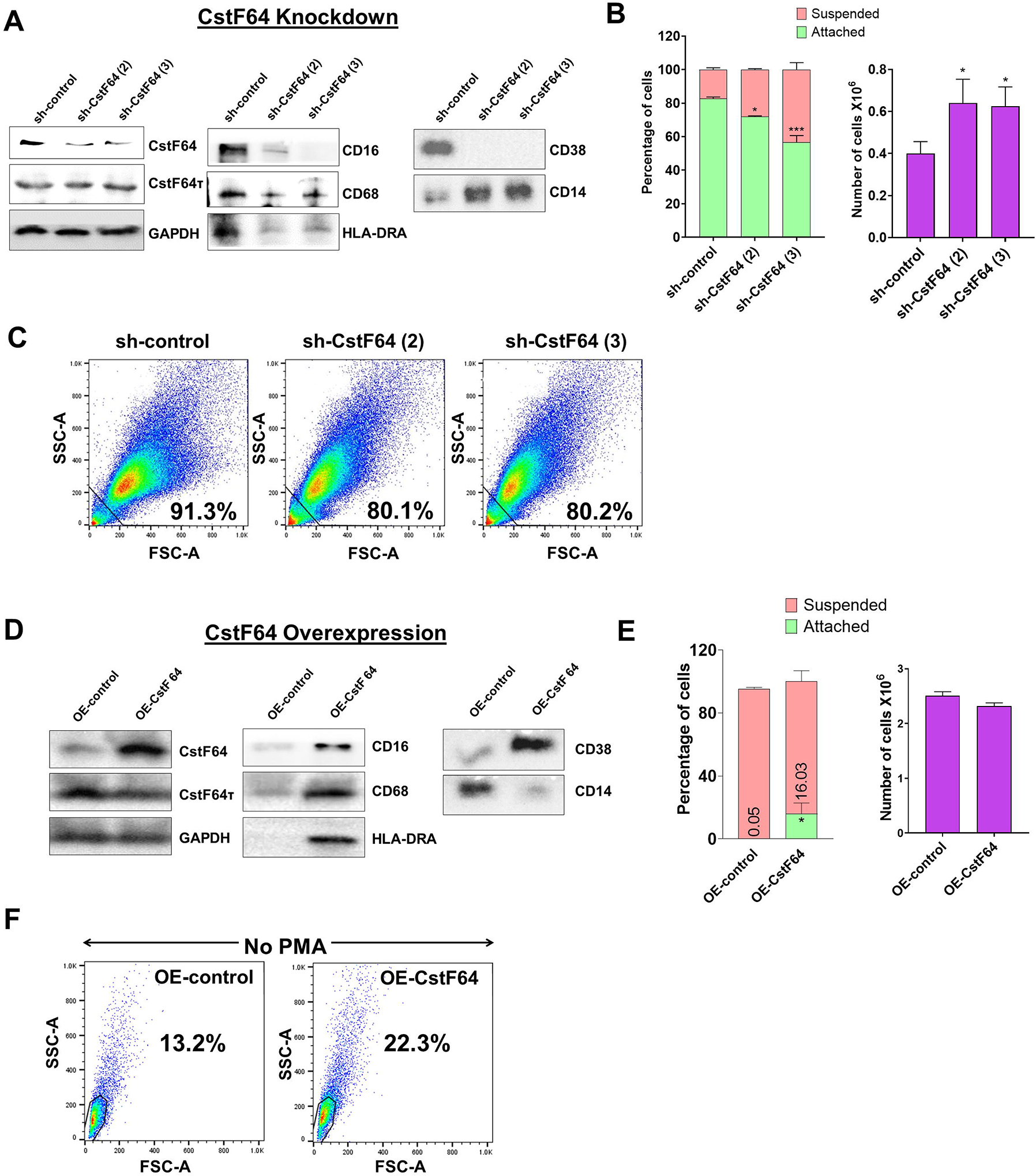
Knockdown or overexpression of CstF64 in U937 cells alters macrophage differentiation. (A) Western blot analysis of C/P factors CstF64 and CstF64τ, macrophage-specific markers CD16, CD68, HLA-DRA, CD38 and the monocyte marker CD14 in U937 cells stably transfected with doxycycline-inducible control shRNA and two different CstF64 shRNAs (shRNA 2 and shRNA 3). Addition of 3 nM doxycycline for 3 days supplemented with 30 nM PMA one day prior to harvesting was done for all three cell lines. (B) Attachment assay (left panel) and changes in total number of PMA-treated cells (right panel) after knockdown of CstF64 compared to the shRNA control. Cells were treated with doxycycline, after which equal number of cells were treated with PMA for 24 h and then counted. The figures are representative of at least three independent experiments. (C) Changes in cellular complexity determined by FACs analysis of side (SSC) and forward scattering (FSC) values for 24h PMA-treated U937 cells stably expressing control shRNA and shRNA against CstF64. (D) Western blot analysis of C/P factors CstF64, CstF64τ and macrophage-specific markers CD68, HLA-DR, CD38 and the monocyte marker CD14 in U937 cells stably transfected with control overexpression vector (OE-control) and those overexpressing CstF64 (OE-CstF64) and treated with 3 nM doxycycline for 3 days prior to harvesting. No PMA treatment has been done. (E) Attachment assay (left panel) and changes in total number of cells (right panel) after overexpression of CstF64 compared to the OE-control. (F) Changes in cellular complexity or granularity determined by FACs analysis of side (SSC) and forward scattering (FSC) values for doxycycline-induced U937 cells transfected with OE-control or OE-CstF64. Percentage indicates a quantitation of cells that have altered granularity.

The changes in expression of monocyte-macrophage markers suggested that CstF64 KD might also affect the cellular changes that occur during macrophage differentiation. Indeed, counting of suspended and attached cells after 24 hours of PMA treatment showed that CstF64 KD in U937 cells significantly reduced cell attachment with both shRNAs (Fig. 7B, left panel). Normally by 24 hours, the cells have stopped dividing but are viable. However, the total cell number at 24 hours was greater in CstF64 KD cells compared to the control, indicating a slowing of the PMA-induced cell cycle arrest (Fig. 7B, right panel). Macrophage differentiation is also characterized by the formation of large, structurally complex cells (Fig. 1) (2), and this change can be detected using flow cytometry to measure the side (SSC) and forward scattering (FSC) of live cells (65). An increase in intensities of both FSC and SSC indicate the formation of cells with complex morphological features (also called increased granularity or complexity) upon PMA-mediated differentiation (Supplementary Fig. S7B). U937 cells depleted of CstF64 demonstrated reduced granularity (Fig. 7C).

Next, to determine if overexpression (OE) of CstF64 would promote macrophage differentiation, we generated stably transfected U937 cells inducibly overexpressing this C/P protein. CstF64 was efficiently overexpressed after 72 hours of doxycycline exposure, while CstF64τ levels did not change (Fig. 7D). The cell proliferation rate was not affected by CstF64 OE (Fig. 7E, right panel). However, overexpression increased expression of macrophage markers such as CD68, HLA-DRA and CD38. and decreased CD14 expression (Fig. 7D). In addition, there was an increase in the number of attached cells from 0 to 16% (Fig. 7E, left panel) and by FACS analysis, U937 cells overexpressing CstF64 demonstrated an almost two-fold increase in granularity (Fig. 7F). It is important to note that these cells were not treated with the differentiation inducer PMA.

In addition to these changes in cellular properties, 3’ RACE analysis showed that CstF64 KD resulted in increased usage of the distal poly(A) sites of the mRNAs *SERPINB1* and *PTCH1,* which would otherwise be shortened after 24 hours of PMA treatment, whereas CstF64 OE without PMA treatment caused decreased usage of the distal poly(A) sites of these mRNAs (Fig. 8). This finding is consistent with studies in other systems showing that CstF64 promotes use of proximal poly(A) sites (55, 66, 67).

**Figure 8.**
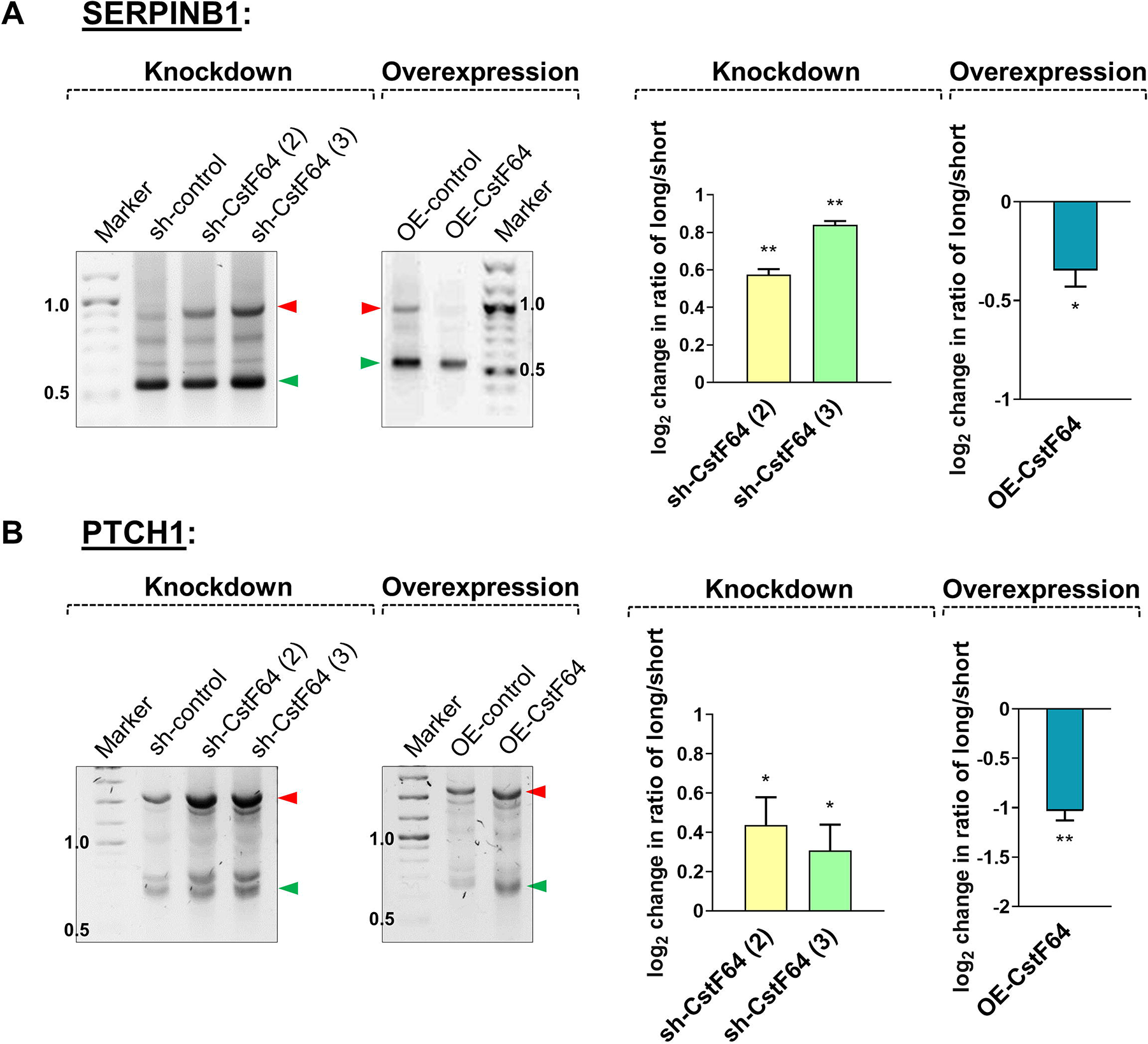
Knockdown or overexpression of CstF64 in U937 cells alters APA. Representative 1% agarose gel images for the 3′ RACE RT-PCR in U937 cells for shortened genes *SERPINB1* (A) and *PTCH1* (B), using gene specific primers at the last exon-exon junction and a common reverse anchor primer. In each subfigure, the effect of depletion (knockdown) of CstF64 and the effect of overexpression of CstF64 are indicated, using the same treatment protocols as described in Fig. 7. Normalized intensities of long and short bands for each PCR were quantified and a long/short ratio determined and normalized to the shRNA or OE control. The corresponding graphs (right) represents mean ± SE from at least two independent experiments. P value <0.05 was considered significant, where * = P ≤ 0.05; ** = P ≤ 0.01; *** = P ≤ 0.001.

In summary, depletion of CstF64 caused changes in cell properties indicative of poorer macrophage differentiation and shifts to longer mRNA isoforms of genes which are normally shortened during differentiation. In contrast, CstF64 overexpression caused the appearance of differentiated phenotypes (cell surface markers, attachment, granularity, and transcript shortening) in the absence of the differentiation inducer.

### 3 Discussion

Macrophage activity is needed for a robust response to infection and tissue damage but when deregulated, it also contributes to many human diseases (3–7). As such, it is critical that monocyte to macrophage differentiation is appropriately regulated to preserve the balance of various monocyte and macrophage populations and ensure the appropriate functional consequences. The findings of our study indicate that regulation of polyadenylation is one of the ways in which gene expression is controlled during macrophage differentiation.

### 3.1 Macrophage differentiation is characterized by both 3′ UTR shortening and lengthening events

APA profiles are known to vary according to the differentiation and growth state of cells, with a general shift to distal poly(A) sites (3’ UTR lengthening) often observed with terminal differentiation and decreased proliferation (12, 57, 68, 69). However, there are counter-examples to this generalization: spermatogenesis and differentiation of secretory cells leads to a prevalence of shorter mRNA isoforms (21, 70), and when B cells are transformed with Epstein-Barr virus, there is only a slight preference for shortening (48). We found that upon macrophage differentiation, the distribution of mRNA isoforms of many genes shifts as the monocyte precursor stops dividing and changes morphology. There were similar numbers of 3’ UTR shortening and lengthening events, indicating that exit from the cell cycle during macrophage differentiation is not associated with predominant lengthening of mRNA isoforms. GSEA analysis showed that lengthened and shortened gene sets are both enriched in groups of genes important for macrophage differentiation and function. We have validated several APA events by 3’RACE in both the U937 and THP-1 monocytic cell lines, and some of the genes examined by 3’RACE have functions relevant to macrophage biology. For example, *SERPINB1* has been reported to influence the innate immune system during influenza infection (71), *PTCH1* is important for macrophage chemotaxis during gastric inflammation (72), and *PSAT1* regulates the inflammatory response in macrophages (73). Our analysis shows that the APA profiles are highly dynamic during macrophage differentiation. Moreover, the APA changes are not globally affecting all genes with multiple poly(A) sites but target sets of genes whose functions are consistent with the need to exit the cell cycle and develop properties of a mature macrophage.

For many genes, 3’ UTR shortening has been shown to increase mRNA and protein expression (44, 74, 75). However, other studies have demonstrated that global 3′-UTR shortening in variety of tissues and cell types has limited influence on mRNA or protein levels (47–49). Similar to these latter studies, our global analysis showed that the direction of APA change is not a good general predictor of mRNA expression levels during macrophage differentiation. It is likely that in this transition, the final levels of different mRNAs are determined by APA-dependent and APA-independent changes in mRNA stability as well as changes in transcription. Moreover, the function of some APA changes during macrophage differentiation may be to affect other mRNA properties such as translatability and localization rather than stability. In summary, our findings support the idea that APA is used as an additional mechanism to insure the timely expression and protein output of critical groups of genes during the transition of monocytes to macrophages.

### 3.2 The increase in C/P proteins may facilitate APA during macrophage differentiation

Our study of the U937 and THP-1 monocytic cell lines shows that differentiating monocytic cells display an overall trend of increased expression of proteins of the core C/P machinery. While there were some differences which could be expected given the different origins of the two lines, the general features of shifts in poly(A) site usage and increase in C/P proteins are consistent between the two. This finding was unexpected, as many studies have shown that the expression of C/P mRNA and protein increases with increased proliferation (56, 66), but monocyte cell division ceases upon induction. The boosted expression of C/P proteins in mature macrophages was accompanied by an increase in the processing efficiency of genes with single poly(A) sites. The specific set of cellular changes accompanying macrophage differentiation, including increased mobility and metabolic activity, may necessitate an increased polyadenylation capacity.

Many studies have demonstrated that the core components of the mRNA 3′ processing machinery are also key regulators of global APA profiles. In macrophages compared to monocytes, we found increases in C/P proteins such as Fip1 and CstF64, known to promote use of upstream sites and production of mRNAs with shorter 3’ UTRs (56, 63, 64, 66, 67). As described below, we showed that manipulation of CstF64 levels indeed altered APA in these cells. The increases in subunits of CFI_m_, which is documented to promote use of downstream sites (74, 76-80) and Pcf11, whose effect can vary depending on gene length and cell type (81–83), could explain why we observe APA changes in both directions.

### 3.3 CstF64 levels alter monocyte-macrophage differentiation and switch poly(A) site choice

In this study, we examined the role of CstF64 in macrophage differentiation. CstF64 is the component of the core C/P complex that recognizes GU/U-rich signal sequences located downstream of the poly(A) site, and it also facilitates APA in numerous cellular contexts. In a well-studied example, an increase in CstF64 protein during terminal B-cell differentiation increases use of an intronic poly(A) site in immunoglobulin heavy chain pre-mRNAs that changes the protein product from a membrane-bound form to a secreted form (26, 63). CstF64 also promotes use of weaker proximal poly(A) sites in proliferating fibroblasts (66), HeLa cells (67) and activated T cells (55), and non-canonical poly(A) site usage (84) as well as processing of specific acetylcholinesterase isoforms (85). Depletion of CstF64 suppressed proliferation of lung cancer cells and exogenous expression promoted proliferation of HEK293 cells (86). CstF64 has also been shown to regulate the cell cycle by controlling replication-dependent histone mRNA 3’ end processing (87, 88), and CstF64 loss blocked the conversion of mouse embryonic stem cells into the endodermal lineage, but not into ectoderm or mesoderm (89). The same study also showed that CstF64 was needed for cardiomyocyte differentiation but not for neuronal differentiation (89), suggesting cell-specific requirements for CstF64. Activation of RAW 264.7 macrophage-like cells with LPS blocked proliferation and caused an increase in CstF64 but not other C/P subunits and an increase in proximal poly(A) site selection (27). As these examples clearly demonstrate, CstF64 is a major player in APA that can promote proliferation, differentiation or specific responses depending on the cell context. However, its role in the differentiation of monocytes to macrophage, a process that involves cell cycle exit and striking morphological changes, has not been previously studied.

In the current work, we found that CstF64 expression increased 6-8 fold during differentiation of monocytic cell lines. Our subsequent analysis showed that CstF64 depletion not only caused lengthening of transcripts that were otherwise shortened during differentiation but also had functional consequences by slowing the acquisition of macrophage properties such as exit from the cell cycle, attachment, marker expression and morphological changes. In contrast, overexpression of CstF64 stimulated macrophage-like changes and enhanced use of promoter proximal sites. Importantly, these effects of extra CstF64 happened in the absence of the powerful inducer used to initiate differentiation. Thus, our findings show that CstF64 is affecting pathways important for macrophage differentiation. They are also in agreement with studies in other systems showing that CstF64 promotes use of promoter proximal sites. However, the increase in other C/P proteins and the observation that some transcripts lengthen during differentiation suggests that CstF64 is not the only driver and that other C/P factors and RNA-binding proteins with roles in APA regulation (12) will also influence isoform expression in macrophages. While our study with CstF64 provides evidence that regulation of polyadenylation is one of the mechanisms that promotes efficient macrophage differentiation, further research is needed to determine the consortium of proteins that regulate APA during this transition in cell state.

In summary, our study has provided new insights into the regulation of gene expression in the differentiation of macrophages. The increase in levels of C/P proteins during differentiation and the fact that manipulation of the C/P protein CstF64 alters the progression of this differentiation supports a model in which changes in 3’ end processing efficiency and alternative polyadenylation are coordinated with changes in transcription and mRNA stability, and this regulatory network helps create mature macrophages poised to respond appropriately to a variety of extracellular stimuli through specific activities such as migration, phagocytosis, antigen presentation, and immunomodulation. Given the critical role of macrophages in regulating the inflammatory response, a greater understanding of how APA promotes beneficial and harmful macrophage activity could open new therapeutic treatment avenues.

## 4 Materials and methods

### 4.1 Bacterial strains, DNA constructs and antibodies

NEB® Stable Competent *E. coli* (Cat. #C3040) were used for transformation of plasmids. All siRNAs and primers were purchased from Eton Biosciences (San Diego, CA). Inducible Lentiviral shRNA constructs targeting CstF64 were purchased from GE Healthcare Dharmacon, Inc. (sequence included in Supplementary Table S1). For overexpression (OE) of CstF64, we cloned CstF64 coding sequence into pLIX_402 vector backbone (a gift from David Root, Broad Institute, MA) and pLIX_402 vector (Addgene plasmid #41394) was used as control. Antibodies were purchased from Santa Cruz, Bethyl and Abcam (Supplementary Table S2).

### 4.2 Cell culture, treatment and microscopy

All cells were maintained in RPMI 1640 medium supplemented with 2 mM L-glutamine and 10% heat-inactivated fetal bovine serum (FBS) at 37°C in a humidified atmosphere of 5% CO_2_. Differentiation of the U937 (ATCC #CRL-1593.2) and THP-1 (ATCC #TIB-202) human monocytic cell-lines into macrophages was performed by treatment with 30 nM phorbol-12-myristate-13-acetate (PMA; Sigma-Aldrich) for up to 24 hours. This PMA concentration has been shown in previous studies to efficiently differentiate U937 cells (30, 90), and as described in the Results section, we found it to be effective with both U937 and THP-1 cells. For monitoring morphological changes, cells were cultured in 6-well plates at a density of 1×10^6^ cells/well, after which they were examined and photographed using an EVOS FL fluorescence phase contrast microscope (Thermo Fisher Scientific). All results were obtained from three independent experiments.

### 4.3 Western blot analysis

Western blot was performed with total cell extracts of untreated and differentiated cells. For preparation of total cell extract, 1X RIPA buffer (20 mM Tris-HCl pH 7.5, 150 mM NaCl, 1 mM Na_2_EDTA, 1 mM EGTA, 1% NP-40, 1% sodium deoxycholate, 2.5 mM sodium pyrophosphate, 1 mM β-glycerophosphate, 1 mM Na_3_VO_4_ and 1 μg/ml leupeptin) was added, followed by incubation on ice for 15 min, centrifugation at 25,000 rcf for 10 minutes at 4°C and supernatant collection. Protein concentration was measured by BCA reagent (Pierce, Thermo Fisher Scientific, Waltham, MA), and 50-80 μg protein was separated on a 10% polyacrylamide-SDS or bis-tris gel and transferred to PVDF membrane. The membrane was blocked for 1 h in 5% non-fat dried milk in TBS-T (Tris Buffer Saline with Tween-20, *i.e.*, 0.05% Tween 20 in 1X TBS) buffer followed by rocking overnight at 4°C with primary antibodies (antibody details provided in Supplementary Table S2). Next day, unbound antibodies were removed by 4 × 5 min washes with TBS-T buffer. The membrane was then incubated with an HRP-conjugated secondary antibody at a dilution of 1:5000 for 1 h at room temperature followed by 3 × 5 minute washes with TBS-T and a final wash with TBS. The blot was developed with SuperSignal™ West Pico PLUS or Femto Chemiluminescent Substrate (Thermo Fisher Scientific) and visualized with a ChemiDoc XRS+ System (Bio-Rad) and quantitated using Image J (91). ACTB or GAPDH was used as the control for individual western blots because their levels did not change upon differentiation. Pre-stained protein markers were used as internal molecular mass standards, and each western blot was performed in three biological replicates of each time point of the differentiation process.

### 4.4 RT-qPCR analysis to quantitate mRNA expression

Total RNA extraction was carried out on 2×10^6^ cells using Trizol according to manufacturer’s protocol. 1.5 μg of RNA was subjected to reverse transcription using oligo dT primer and Superscript III reverse transcriptase. The cDNA was amplified by qPCR with specific primers (Supplementary Table S1) using the C1000™ thermal cycler with CFX96 Touch Real-Time PCR Detection System (Bio-Rad Laboratories). The relative expression of genes was analyzed quantitatively by the ΔΔC_t_ method. ACTB RNA was used as the normalization control for RT-qPCR-based RNA expression analyses because its level did not change upon differentiation according to the poly(A)-seq datasets. Primers for RNA expression were designed at the first exon according to the annotation featured in Ensembl (92).

### 4.5 RT-qPCR analysis for quantitating processing activity

We used a previously published assay to measure the processing efficiency at the single poly(A) sites of the MYC and ACTB genes (60, 61). This assay uses random hexamers for reverse transcription of 1 μg of total RNA. RT-qPCR is then performed with a primer pair upstream of the poly(A) site, called “US” which measures the level of unspliced transcripts and another primer pair that spans the poly(A) site, called “span”. The sequences of these primers are provided in Supplementary Table S1.

### 4.6 Poly(A)-Click sequencing and analysis

This was a paid service from Click-Seq Technologies, in which total RNAs isolated from control and differentiated U937 cells were subjected to Poly(A) Click-sequencing as described (39, 40, 43). Briefly, reverse-transcription with oligo-dT primer (unanchored) supplemented with small amounts of spiked-in 3’-azido-nucleotides (AzVTPs) was performed to initiate cDNA synthesis from within the poly(A) tail, ensuring stochastic termination upstream of the poly(A) tail. The resulting 3’-azido-blocked cDNA molecules were purified, ‘click-ligated’ and amplified to generate a cDNA library. These libraries were pooled and sequenced using the manufacturer’s standard operating procedures on a NextSeq 550, obtaining 1×150 bp (130M) paired-end reads. Raw data from the sequencing reactions for four biological replicates for 0h vs 24h and two biological replicates for 0h vs 6h was analyzed by Click-Seq Technologies using Differential Poly(A)-Clustering (DPAC), a pipeline to pre-process raw poly(A)-tail targeted RNAseq data, map to a reference genome, identify and annotate the location of poly(A) sites, generate poly(A)-clusters (PACs) and determine the differential abundance of PACs and differential gene expression between two conditions (40). The raw reads and mapped reads for each library are provided in Supplementary Table S2 (sheets 1 and 2). Each of these libraries were trimmed and quality filtered as described (43). Gene counts collapse all mapped poly(A) sites within that one gene into one count, this provides differential gene expression data. On average, 87% of all the mapped reads were found within or 200 nucleotides downstream of annotated mRNAs. Multiple poly(A) sites occurring within 25 nucleotides of one another were grouped into a poly(A) cluster (PAC) and treated as a unit to determine alternative polyadenylation events within exons and introns. The cutoff to be considered a PAC is 10 counts. In the final stage of DPAC, the mapped reads from each individual samples are used to determine the frequency of PACs in each dataset by determining the overlaps of the 3’ ends of the mapped reads with the provided poly(A) cluster database using bedtools (93). Data normalization and statistical tests are applied using the canonical DESeq2 (94) pipeline using local dispersion estimation and Independent Hypothesis Weighting (IHW) (95) to estimate false discovery rates and power maximization. The final output of DPAC for the 6h and 24h PMA-treated U937 samples are provided in Supplemental Tables S4 and S5.

Principal component analysis (PCA) comparing control and PMA-treated U937 cells (0h vs 6h and 0h vs 24h) was done with the help of the DESeq2 pipeline (94) to determine and identify outlying samples within datasets. Variables (i.e. the times of differentiation) are transformed into orthogonal principal components where each principal component indicates the direction with the greatest variability within the data. PCA plots (Supplementary Fig. S8A, B) revealed that treatment time was the most critical variable. Sequencing data sets are uploaded to NIH-GEO with the accession number #GSE169140. Volcano plots illustrating changes in gene counts and PAC counts (Supplementary Fig. S8C, D) were generated for both 6h and 24h PMA treated U937 cells by plotting the log_2_ fold change values vs p_Adj_ value. Differential Gene or PAC usage is defined as >1.5 fold change with p_Adj_ <0.1. The volcano plots indicate that the number of differentially expressed genes and the number of PACs showing change in usage increases as differentiation progresses. Gene Set Enrichment Analysis (GSEA) was carried out using online resources from the Broad Institute (50). Poly(A) site locations for genes were extracted into BED files and visualized with the help of UCSC genome browser (96).

### 4.7 3’ RACE

We modified the previously published 3’RACE protocols for this study (97, 98). Briefly, total RNA extraction was done using Trizol according to manufacturer’s protocol. 1 μg of RNA was subjected to reverse transcription for 1 hour using 100 units of Superscript IV reverse transcriptase and an oligo(dT)-containing adapter primer to poly(A) mRNA. After cDNA synthesis, the RNA template is removed from the cDNA:RNA hybrid molecule by digestion with 2 units of RNase H after cDNA synthesis to increase the sensitivity of PCR. For purification, the prepared cDNA was then chloroform extracted and precipitated in 0.3M sodium acetate and 3 volumes of ethanol. Subsequently, cDNA was subjected to PCR reaction with HotStarTaq Master Mix Kit (Qiagen) utilizing gene-specific forward primer (designed at the junction of the last two exons) and a unique anchor primer complimentary to the oligo(dT)-containing adapter primer (sequences provided in Supplementary Table S1). The first primer extension was accomplished by incubation at annealing temperature for 1 min, followed by incubation at 72°C for 40 min. Exponential amplification was accomplished by 30-35 cycles of PCR at 94°C for 1 min, 55°C /60°C (3-5 degrees lower than the lowest primer T_m_) for 30 sec and 72°C for 5 min, followed by an additional 10-min extension at 72°C. The products were resolved in 1% agarose gels along with 1 kb Plus DNA Ladder (New England Biolabs) and visualized using SYBR™ Safe DNA Gel Stain (Thermo Fisher) with a ChemiDoc XRS+ System (Bio-Rad) and quantitated using Image J (91).

### 4.8 Lentivirus construction and cell treatment

For shRNA-based KD, three different lentiviral SMARTvector constructs, designed for doxycycline-inducible expression of shRNA for human CstF64 gene silencing, were purchased from GE Healthcare Dharmacon, Inc. These vectors incorporate the Tet-On® 3G bipartite system to inducibly control of shRNA with the Tet 3G promoter (PTRE3G), while the expression of the Tet-On® 3G transactivator protein is under the control of a constitutive human EF-1 promoter for optimal expression in the monocyte-macrophage system. A SMART vector inducible lentiviral control shRNA was used to control for potential non-specific effects caused by expression of shRNA. The sequences of the shRNAs are in Supplementary Table S1. As shown in the Results, shRNA #2 & 3 gave efficient KD but shRNA #1 was not effective. To generate stable cells inducibly overexpressing CstF64, its coding sequence was amplified by PCR from U937 cDNA and cloned into the pLIX_402 inducible lentiviral expression vector by using NheI and AgeI restriction enzyme sites to generate the OE construct. The primers used for cloning are mentioned in Supplementary Table S1.

These recombinant plasmids were co-transfected with the components of Dharmacon™ Trans-Lentiviral packaging kit into HEK293FT cells using FuGENE® HD (Promega Corp.) according to manufacturer’s protocol. Transfection of HEK293FT cells was done in 6-well plates when the cells were 80-85% confluent, and transfection media changed after 16 hours. The recombinant lentiviruses were harvested at 48 h post-transfection, spun at 1250 rpm for 5 minutes and filtered by a 0.45 μm filter to remove cells and debris. Purified virus was used for infecting monocytic cells. 1.5×10^6^ target cells were seeded in 4 ml of media per well in a 6-well plate and cultured overnight. Lentiviral particles (1-2 ml per well) were added the next day to the cells in culture media containing 10 μg/ml polybrene for efficient infection. Selection of cells stably expressing CstF64-shRNAs or control-shRNA and pLIX_402 overexpressing CstF64 or control pLIX_402 started 72 h post-transfection or a time determined by the expression of GFP from the cells. Growth medium was replaced with fresh selection medium containing 1 μg/mL of puromycin. Puromycin-containing medium was refreshed every 2–3 days, and selection was completed after approximately 1 week, after which clones were expanded for 2 more weeks and frozen for later use. The Tet-On 3G induction system was activated by 3 nM doxycycline for 3 days for both KD and OE transfections. Additionally for KD clones, 30 nM PMA was added one day prior to harvesting for RNA and protein extraction.

### 4.9 Attachment assay

U937 and THP-1 cells were either untreated or treated with PMA and left for 24 hours to attach. After that, unbound cells were collected from the plate by washing gently with PBS and collected for counting (denoted as “suspended”). The attached cells were collected by trypsinizing to release them from the plate. Cells were then stained with trypan blue and counted in a hemocytometer and represented as percentage of suspended or attached cells over total (suspended + attached). The figures are average of three independent observations.

### 4.10 Flow cytometry

Phenotypic analysis of monocytes and macrophages was performed using flow cytometry. After differentiation, cells were recovered from culture plates and washed with phosphate-buffered saline (PBS) containing 1% paraformaldehyde for fixation. In order to avoid non-specific binding and background fluorescence of cells expressing high levels of Fc receptors, the cells were incubated with Fc Receptor Blocking Solution (Human TruStain FcX, Biolegend) according to the manufacturer’s instructions for 10 minutes, followed by washing with cold PBS. Zombie Aqua™ Fixable Viability kit (BioLegend) at a final dilution of 1:2000 was used for the exclusion of dead cells. Thereafter, cells were analyzed with a LSR II flow cytometer (BD Biosciences) using FACSDiva software, and data were analyzed with the FlowJo software. Primary gating was done on the basis of forward versus side scatter (FSC vs SSC) to compare size and granularity (complexity) of live cells across samples (31).

### 4.11 Statistical analysis

All experiments were performed in at least three independent sets. Data are presented as mean ± SE. Statistical analysis was performed using GraphPad Prism 6.01 (GraphPad Software Inc., La Jolla, CA, USA). Two-way analysis of variance (ANOVA) was performed to determine the significance between the groups. Considerations were * = P ≤ 0.05; ** = P ≤ 0.01; *** = P ≤ 0.001; **** = P ≤ 0.0001. A P value <0.05 was considered significant.

### 4.12 Materials availability

All plasmids and cell lines will be available upon request.

## Abbreviations

CstF: Cleavage Stimulation Factor
CPSF: Cleavage and Polyadenylation Specificity Factor
C/P: Cleavage and Polyadenylation
KD: Knockdown
OE: Overexpression
UTR: Untranslated Region

## 6 Author Contributions

CM and SM conceived the study and CM provided general oversight. SM developed the strategy and methodology, acquired the data, reported and organized the findings. CM and SM designed the experiments, interpreted the results, and wrote and revised the manuscript. JG contributed to the computational analyses.

## 8 Acknowledgments

We thank Tufts University faculty Alexei Degterev and Karl Munger for providing cell lines and Marta Rodriguez-Garcia for consulting with us on FACS analysis. We thank members of Karl Munger’s lab for use of the cell culture facility and want to acknowledge the critical inputs of the members of FACS core, Tufts University, and other members of the Moore lab. We would also like to thank Andrew Routh and Elizabeth Jaworski of ClickSeq Technologies for their valuable input on the bioinformatic analysis, and Nathaniel Maki and Chris Wilson of Mount Desert Island Biological Laboratory for helping perform the alignments for RNA-seq analysis.

## 10 Data Availability Statement

Poly(A) sequencing data sets are available in the NCBI Gene Expression Omnibus (GEO), accession number #GSE169140. Supplementary materials are attached with this manuscript.

