## Supplementary figures and tables for "Macrophage differentiation is marked by increased abundance of the mRNA 3’ end processing machinery, altered poly(A) site usage, and sensitivity to the level of CstF64": Supplementary material SM.pdf

### 1 Supplementary Figures and Tables

#### 1.1 Supplementary Tables

**Supplementary Table S1. Primers and shRNA sequence used in this study.**

| Purpose | Primer name | Forward primer 5'-3' | Reverse primer 5'-3' |
|---|---|---|---|
| RT-qPCR for gene expression | CD86 total mRNA | GACTGAGTAACATTCTCTTTGTGATGGCC | CTACTAGCTCACTCAGGCTTTGG |
| | ICAMI total mRNA | CCAATGTGCTATTCAAACTGC | CAGCGTAGGGTAAGGTTCTTG |
| | JUN total mRNA | AAGTTGCGACGGAGAGAAAA | ACGCAACCCAGTCCAACTT |
| | FOS total mRNA | GACCGAGCCCTTTGATGAC | GCCACTGTGCAGAGGCTC |
| | NFKB1 total mRNA | CAAGCAGCTCTGCAGCAG | ACTGTCATAGATGGCGTCTG |
| | IKZFI total mRNA | CACGTGATGGACCAAGC | CGGCTTGTGCAGCTGGTAC |
| | OSM total mRNA | CTCCTGGACCCCTATATACG | GTGTGGCATTGAGGGTCTG |
| | IL12A total mRNA | GAAAGTCCTGCCGCGCC | GTTTCTGGCCAAACTGAGGTG |
| | GAPDH mRNA | TGATGACATCAAGAAGGTGGTGAAG | TCCTTGGAGGCCATGTGGGCCAT |
| | ACTB mRNA | CATGTACGTTGCTATCCAGGC | CTCCTTAATGTCACGCACGAT |
| Processing activity | MYC Total | ATTACAGGTGTGAGCCAGGG | AGCCTGCCTCTTTTCCACA |
| | MYC Span | ATCATTGAGCCAAATCTTAAGTTGTG | CTCTGAAGGGGCAATTGATGA |
| | ACTB Total | TCAAGGTGGGTGTCTTTCCT | CCTGCTTGCTGATCCACATC |
| | ACTB Span | GCTTTTGGTCTCCCTGGGA | CTGCACTCTGGGTAAGGACA |
| CstF64 overexpression (For PCR cloning) | | CTTACGGCTAGCGCCACCATGGCGGGTTTGAC (Nhe I enzyme site) | CATTAGACCGGTAGGTGCTCCAGTGG (Ase I enzyme site) |

| Purpose | Sequence |
|---|---|
| CstF64 knockdown shRNA #1 | CGTAGAGAACGATCCACCG |
| CstF64 knockdown shRNA #2 | TGGCATTCCAACCTGAGCC |
| CstF64 knockdown shRNA #3 | CGTGCATCTATTCCTCGGG |
| Oligo d(T) adapter for 3'RACE | GCGAGCACAGAATTAATACGACTCACTATAGGTTTTTTTTTTTTTTVN |
| Reverse anchor-primer for 3'RACE | GCGAGCACAGAATTAATACGACT |
| SERPINB1 3'RACE exon Fwd | GGCCTGAAGAAGATTGAGGAAC |
| PTCH1 3'RACE exon Fwd | CAGCACCTTGTTACAGACAAGAAG |
| PSAT1 3'RACE exon Fwd | CATAGGTCTGTGGGAGGC |
| MRPL44 3'RACE exon Fwd | GCACTTTTCATCAGGGACTTCTTAATTAC |
| CIAPIN1 3'RACE exon Fwd | GTGGAAACTGCTACCTGGGC |
| EIF1 3'RACE exon Fwd | GCGTTTAAGAAAAAGTTTGCCTG |

**Supplementary Table S2. Antibodies used in this study.**

| Catalogue no. | Name | Host | MW |
|---|---|---|---|
| SC-20052 | CD16 Or FCGR3B | Mouse monoclonal | 50 kDa |
| A6554 | CD68 | Rabbit polyclonal | 110-140 kDa |
| SC-55593 | HLA-DRα | Mouse monoclonal | 36 kDa |
| SC-18853 | ICAM1 (6.5B5) | Mouse monoclonal | 85-110 kDa |
| SC-374650 | CD38 (H-11) | Mouse monoclonal | 45 kDa |
| SC -58951 | CD14 (5A3B11B5) | Mouse monoclonal | 53-55 kDa |
| ab8227 (Abcam) | Beta Actin | Rabbit polyclonal | 43 kDa |
| SC-365062 | GAPDH (G-9) | Mouse monoclonal | 37 kDa |
| Bethyl A301-357A | $CFI_m68$ or CPSF6 | Rabbit | 68 kDa |
| Bethyl | $CFI_m25$ or NUDT21 | Rabbit | 25 kDa |
| SC-393880 | $CFI_m59$ Or CPSF7 | Mouse monoclonal | 52 kDa |
| SC-398392 | FIP1L1 (C-10) | Mouse monoclonal | 67 kDa |
| SC-166281 | CPSF160 or CPSF1 (G-10) | Mouse monoclonal | 160 kDa |
| SC-165983 | CPSF100 or CPSF2 (A-11) | Mouse monoclonal | 100 kDa |
| SC-374466 | WDR33 (D-1) | Mouse monoclonal | 146 kDa |
| SC-393001 | CPSF73 or CPSF3 (C-3) | Mouse monoclonal | 73 kDa |
| SC-393316 | CPSF30 or CPSF4 (D-1) | Mouse monoclonal | 30 kDa |
| Bethyl A301-250A | CSTF50 or CSTF1 | Rabbit polyclonal | 50 kDa |
| SC-376575 | CstF-77 (G-5) or CSTF3 | Mouse monoclonal | 77 kDa |
| SC-398862 | CstF64 (H-1) or CSTF2 | Mouse monoclonal | 64 kDa |
| Bethyl A301-487A | CSTF2T/TauCSTF64/CstF64τ | Rabbit polyclonal | 70 kDa |
| Bethyl AAS02642C | hClp1 | Rabbit polyclonal | 41 kDa |
| Bethyl A303-706A | Pcf11 | Rabbit Polyclonal | 173 kDa |
| SC-135410 | Symplekin (H308) | Rabbit polyclonal | 150 kDa |
| SC-32915 | PAP (H-300) | Rabbit polyclonal | 64 kDa (may be 90 kDa) |

#### 1.2 Supplementary Figures

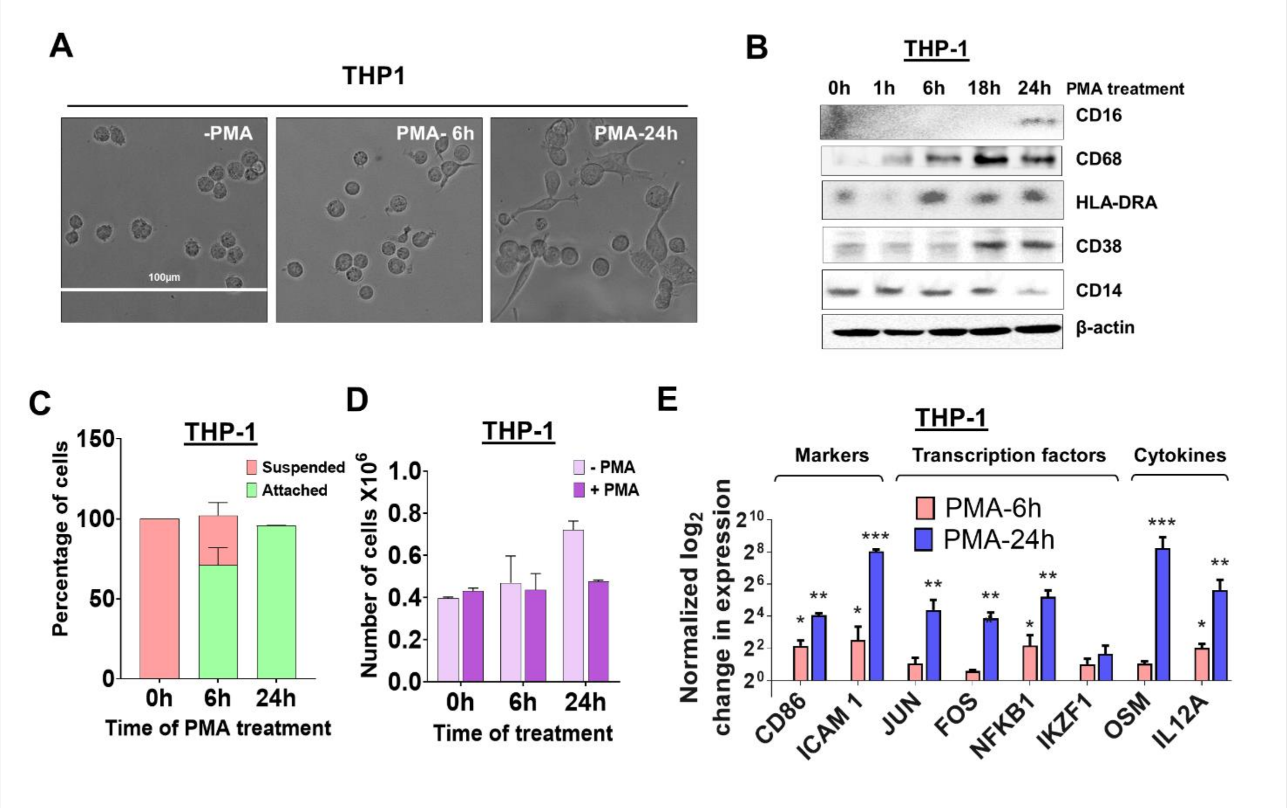


**Supplementary Figure S1. Morphological changes and altered expression of cell-surface markers and mRNAs during differentiation of THP-1 cells and human primary monocytes.**

(A) The effect of differentiation on THP-1 cellular morphology. THP-1 cells exposed to PMA (30 nM) for 6 and 24 hours and observed by phase contrast microscopy (40X). (B) Western blots of total cell lysates from THP-1 cells differentiated with PMA (30 nM) for 0, 1, 6, 18 and 24 hours. Blots were probed with antibodies against CD16, CD68, HLA-DRA, ICAM1, CD38 and CD14 with β-actin as the loading control. (C) Attachment assay for THP-1 cells, performed as described for Fig. 1C. (D) Proliferation assay for THP-1 cells performed as described for Fig. 1D. (E) Effect of THP-1 differentiation on the expression level of mRNAs from genes involved in macrophage differentiation and function. RT-qPCR assays performed for genes as described in Fig. 1E.

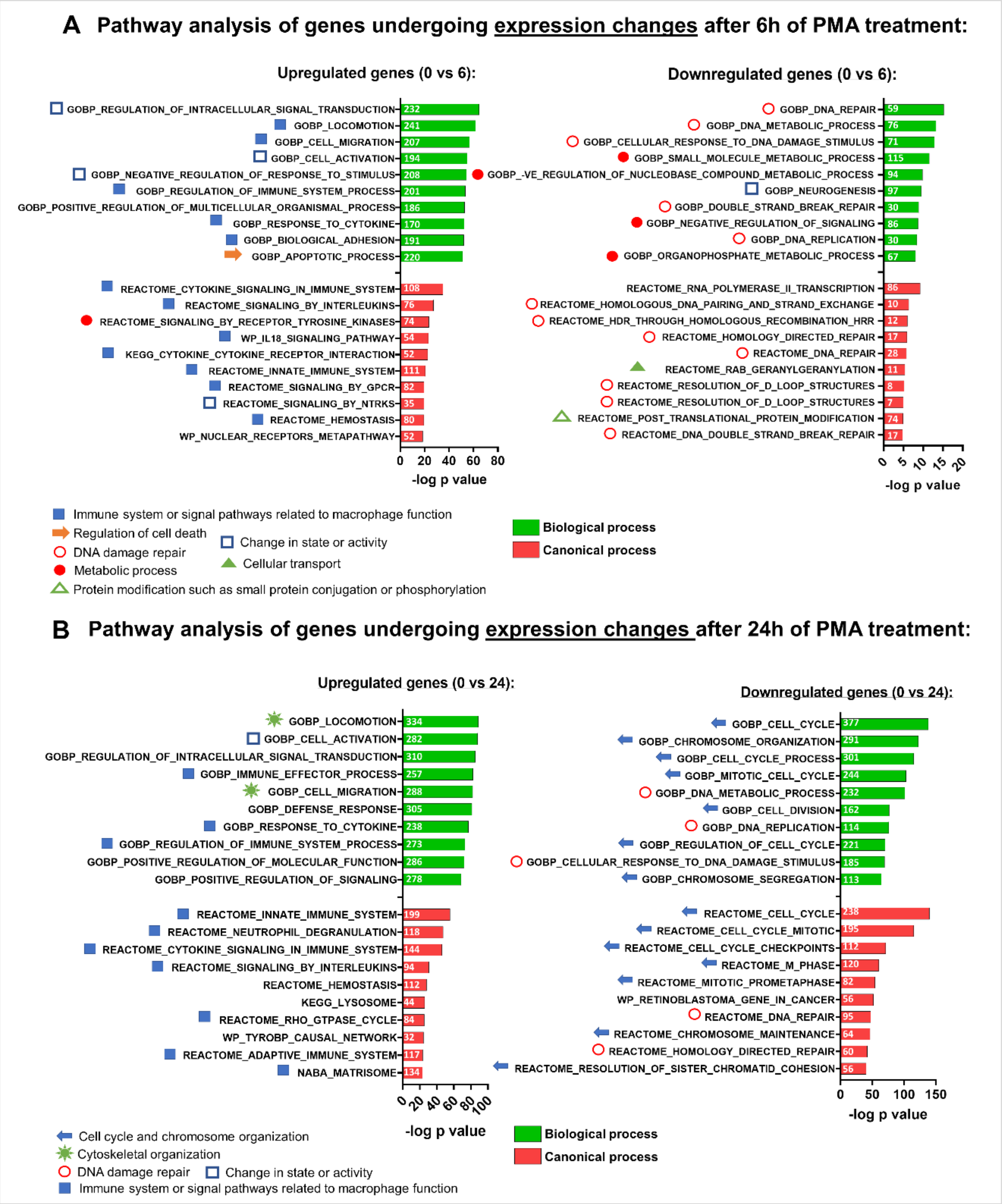


**Supplementary Figure S2. Gene set enrichment analysis of mRNAs with expression changes in U937 cells after PMA treatment.**

Functional annotation clustering of mRNAs whose expression is regulated after 6h (A) and 24h (B) of differentiation. The 10 most significant GO-process enriched gene groups according to GSEA analysis are ranked based on negative $\log_{10}$ (P-values) for upregulated (left panel) and downregulated (right panel) transcripts. Green indicates biological process; red, canonical process. Pathways classified according to specific cellular processes are grouped together as indicated by the individual symbols at the bottom of the graphs.

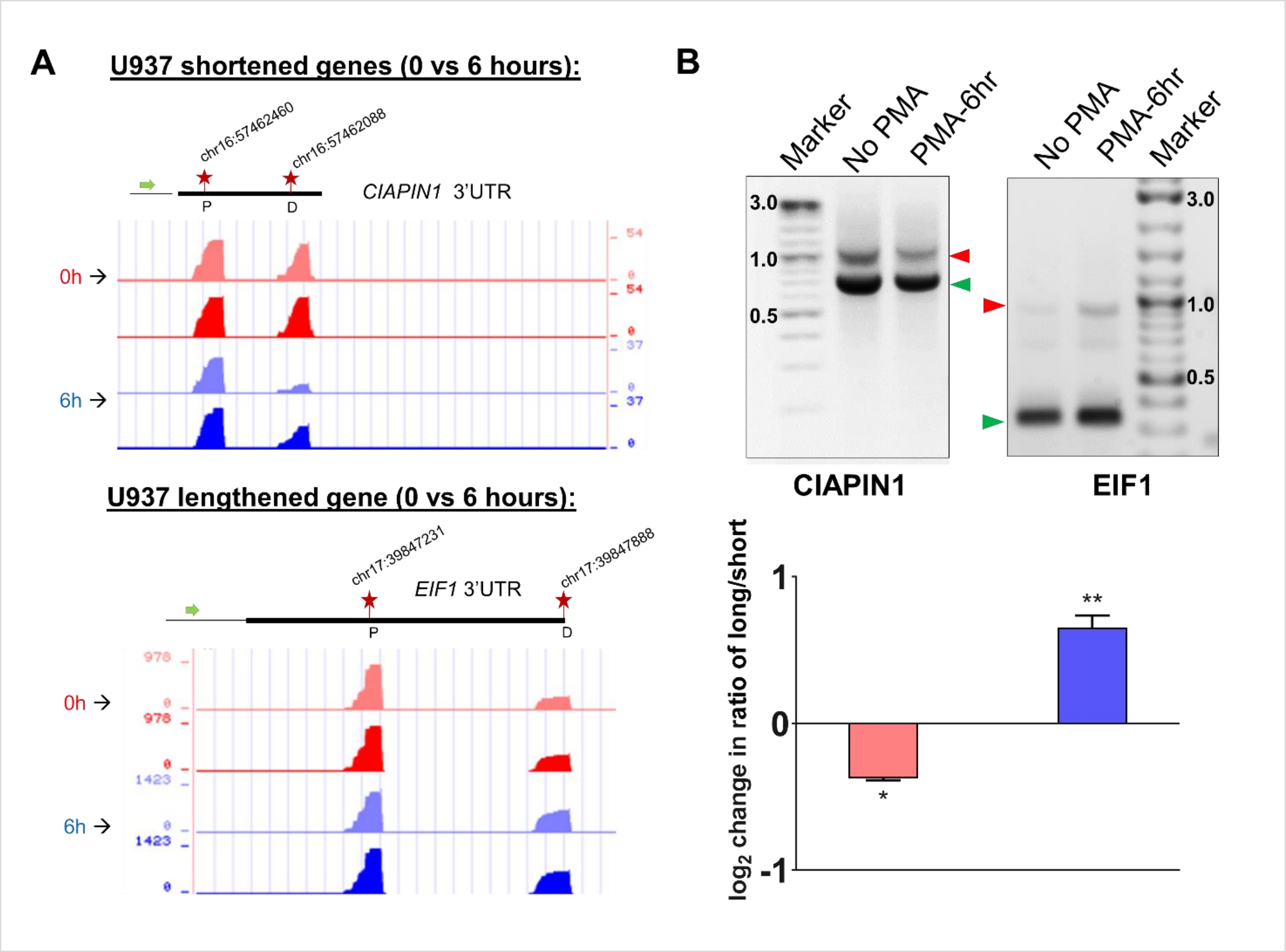


**Supplementary Figure S3. Changes in poly(A) site use in U937 cells after 6 hours of PMA treatment and validation by 3'RACE.**

(A) UCSC genome browser plots of RNA sequencing tracks highlighting the 3′-UTR profile differences for shortened and lengthened genes after 6h PMA treatment with respect to control (0h). The colors of the tracks represent 0h (red) and 6h (blue). Proximal (P) and distal (D) poly(A) sites are indicated with red stars. The green arrow defines the direction of the coding strand, blue arrow defines the direction of chromosome co-ordinates and tag counts are indicated on the y axis. Additionally, positions and chromosome co-ordinates of the annotated PACs are indicated at the top of each browser plot. (B) Representative 1% agarose gel images for the 3′ RACE RT-PCR from U937 cells (untreated and 6h PMA-treated) for shortened gene CIAPIN1 and lengthened gene EIF1 (upper panel) using gene specific primer at the last exon-exon junction and reverse anchor primer. Normalized intensities of long and short bands for each PCR were quantified and a long/short ratio determined and normalized to the No PMA control. The graph (lower panel) represents mean ± SE from at least two independent experiments. P value <0.05 was considered significant, where * = $P \leq 0.05$; ** = $P \leq 0.01$; *** = $P \leq 0.001$.

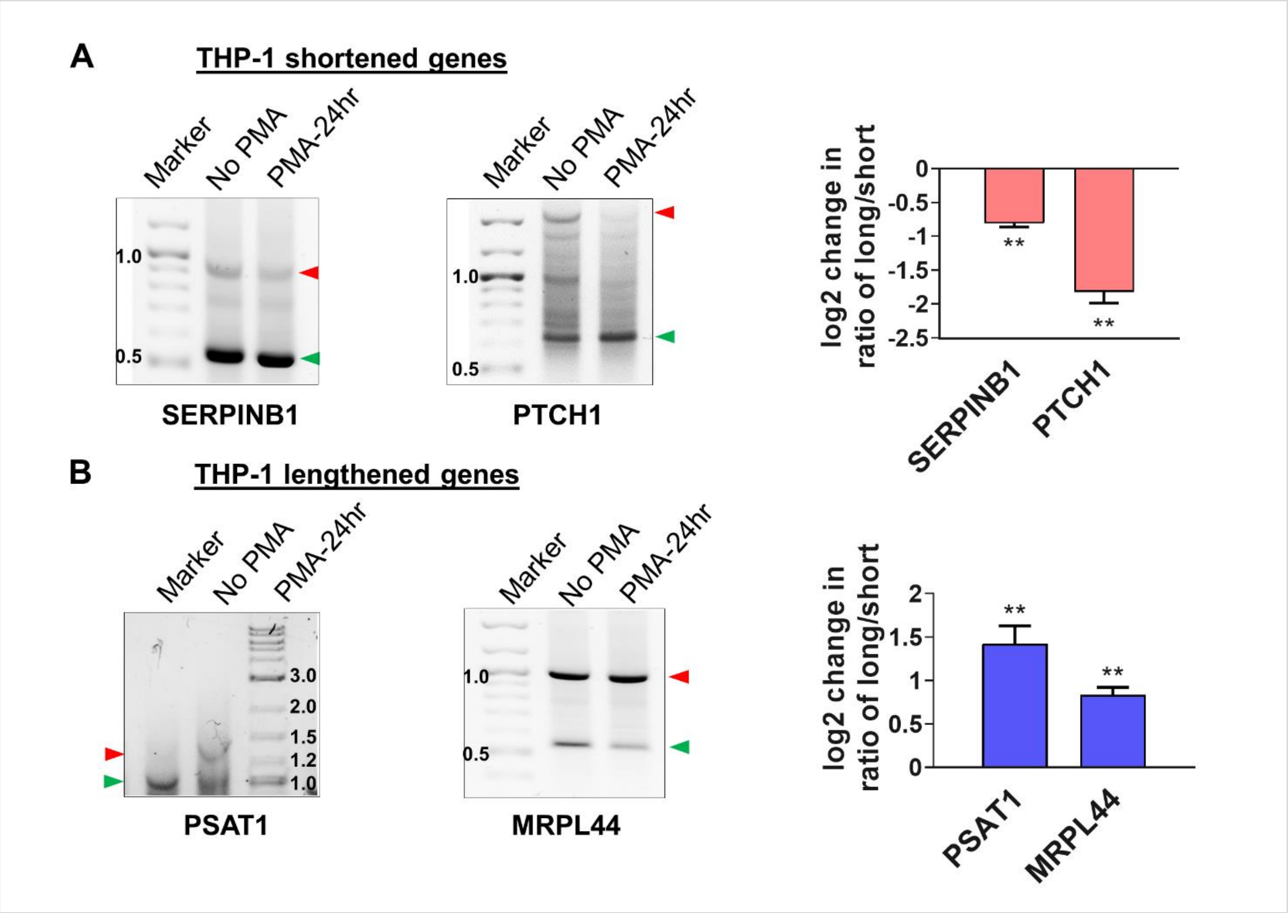


**Supplementary Figure S4. Validation of APA events in THP-1 and human primary monocytes by 3’ RACE.**

Representative 1% agarose gel images for the 3′ RACE RT-PCR from control and differentiated THP-1 cells and human primary monocytes for shortened genes SERPINB1 and PTCH1 (A and C), lengthened genes PSAT1 and MRPL44 (B and D) with gene specific primer at the last exon-exon junction and reverse anchor primer. Normalized intensities of long and short bands for each PCR were quantified and a long/short ratio determined and normalized to the No PMA or undifferentiated controls. The corresponding graphs (right) represents mean ± SE from at least two independent experiments. P value <0.05 was considered significant, where * = $P \leq 0.05$; ** = $P \leq 0.01$; *** = $P \leq 0.001$.

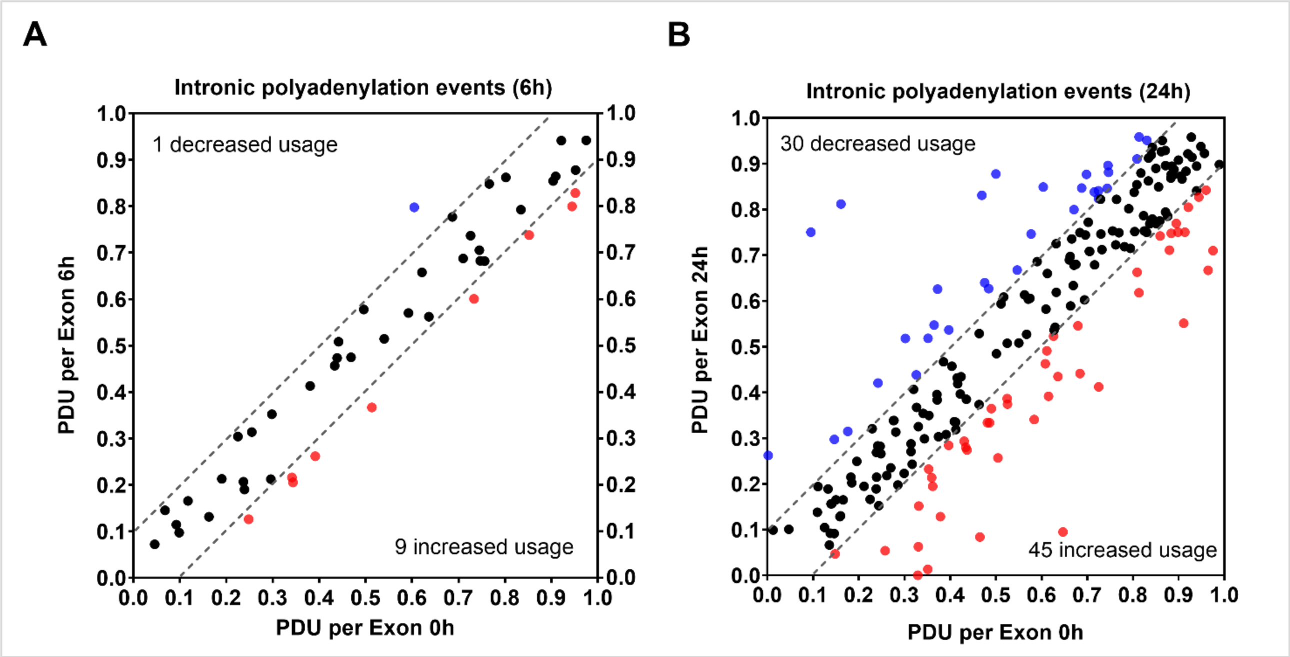


**Supplementary Figure S5. Evidence of intronic polyadenylation events.**

(A) and (B) mRNAs undergoing intronic polyadenylation in U937 cells treated with PMA for 6h and 24h, classified according to their delta PDUI changes in both directions. Every exon with multiple PACs having “splicing APA” event (Supplementary Tables S3 and S4) is plotted here, and the PDU for 0h is on the x-axis and 24h on the y-axis. Depending upon the significance levels reported by the DPAC pipeline, blue dots indicate lengthening or decreased usage of the poly(A) site whereas pink dots indicate shortening or increased usage of the poly(A) site. Dotted grey lines indicate a change in PDU of 0.1 in either direction and the events (black dots) within the lines are considered not significantly altered.

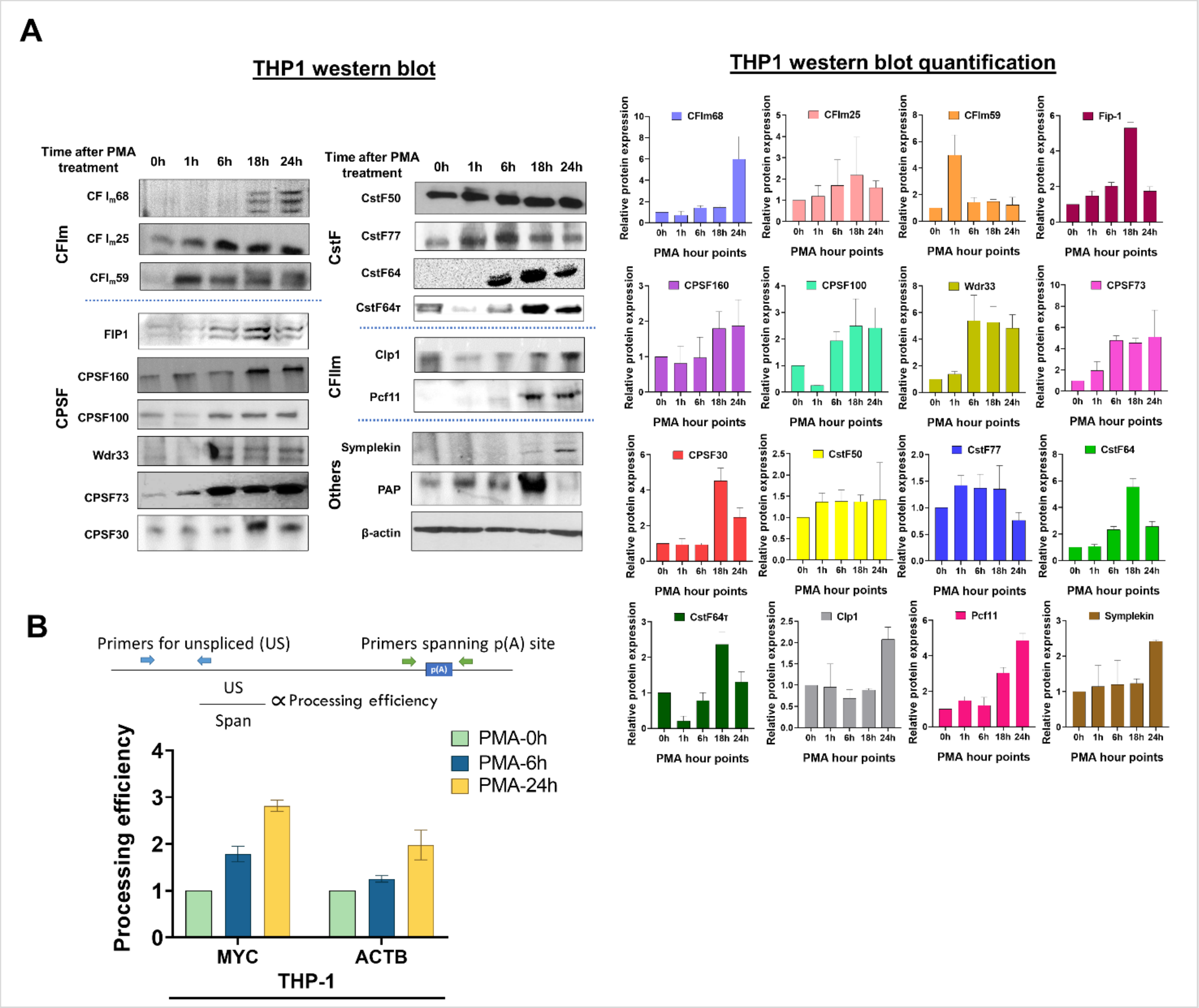


**Supplementary Figure S6. Effect of macrophage differentiation on the expression of C/P proteins and mRNAs in THP-1 cells and human primary monocytes.**

(A) Whole cell lysates from THP-1 cells treated with PMA for 0, 1, 6, 18 and 24 hours were separated by 10% SDS-PAGE and western blotting was performed for the indicated subunits of the C/P complex, with β-actin as the loading control. Each western blot (left panel) was performed in three biological replicates of the differentiation process and the quantified data is shown in the right panel. (B) Processing efficiency of MYC and ACTB transcripts for THP-1 cells. The bar graph shows the ratio of unspliced (US) RNA transcripts (detected by RT-qPCR with a primer pair upstream of the poly(A) site) to unprocessed RNA (detected by RT-qPCR with a primer pair that spans the poly(A) site) as shown in the schematic. The cDNA preparation was done by using random hexamers.

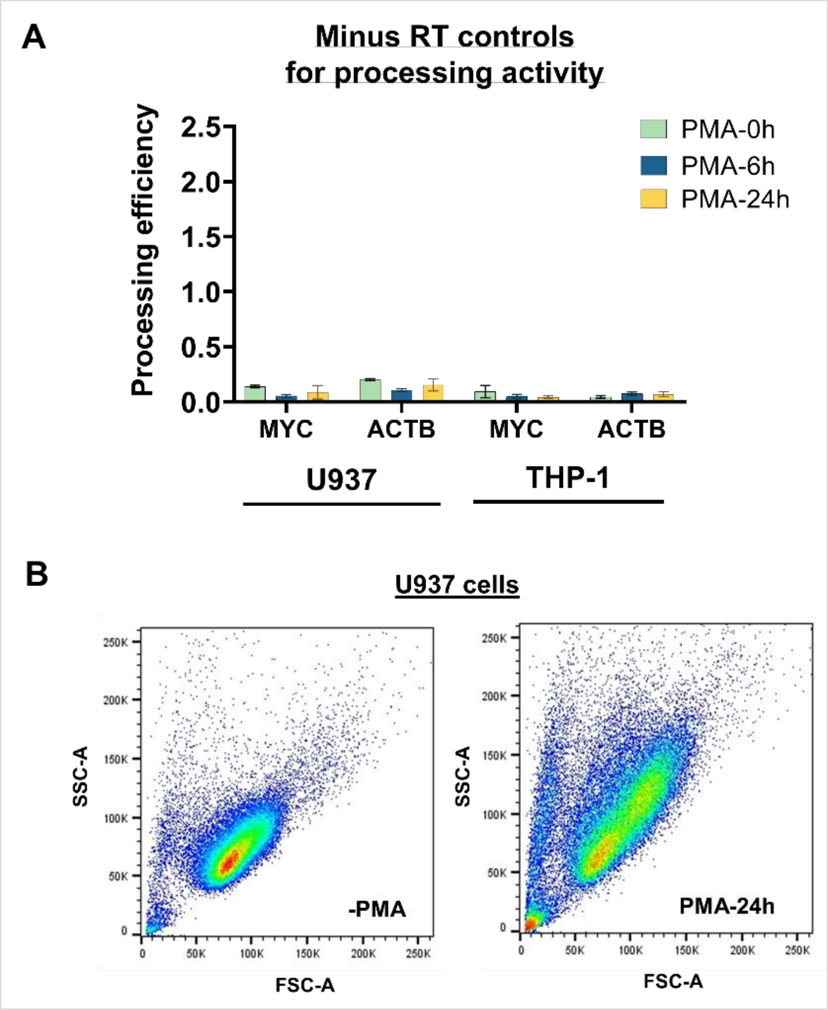


**Supplementary Figure S7. Processing efficiency minus RT control and cellular complexity of U937 cells by FACS.**

(A) Minus RT control for processing efficiency of MYC and ACTB transcripts for U937 and THP-1 cells. PCR was done by adding equivalent amount of RNA that was used to prepare cDNA for Fig. 6D and S6D (1µg). The bar graph shows the ratio of unspliced (US) RNA transcripts (detected by RT-qPCR with a primer pair upstream of the poly(A) site) to unprocessed RNA (detected by RT-qPCR with a primer pair that spans the poly(A) site) as shown in Fig. 6D. (B) Changes in cellular complexity determined by FACs analysis of side (SSC) and forward scattering (FSC) values for untreated and 24h PMA-treated U937 cells.

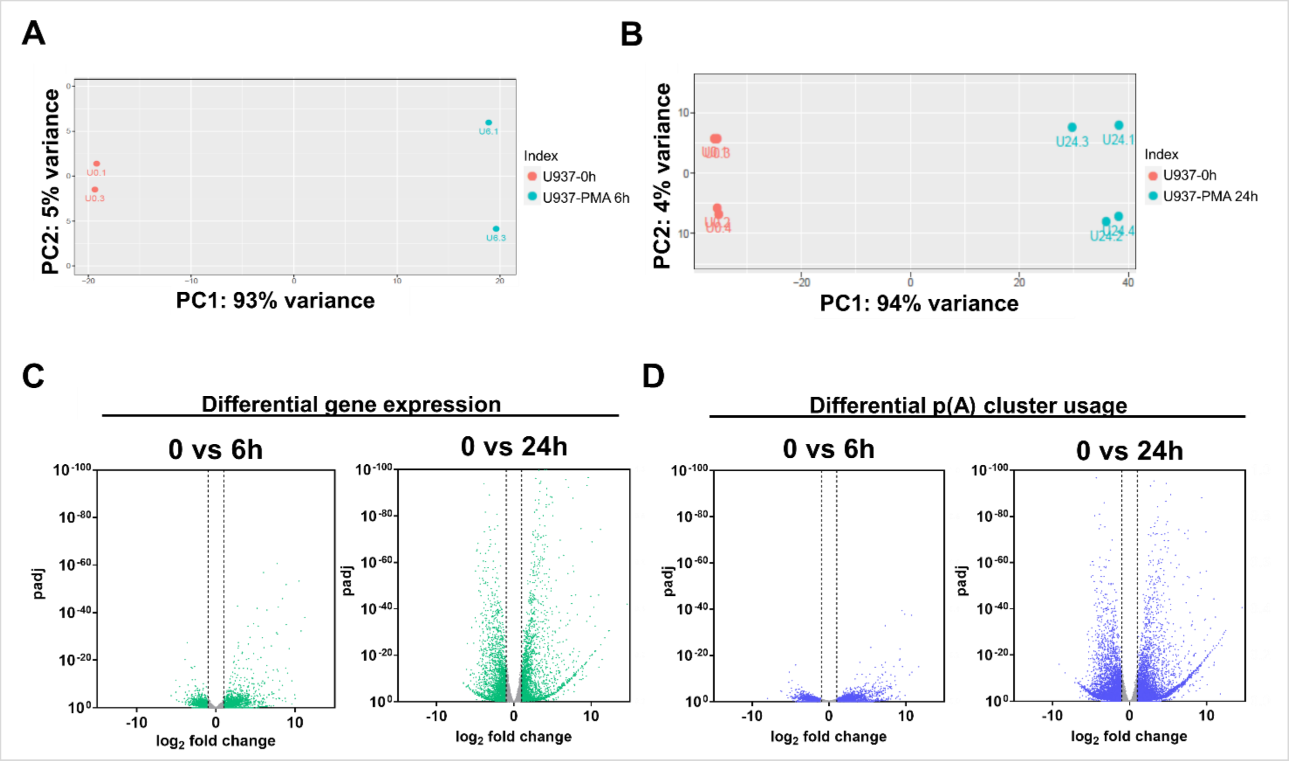


**Supplementary Figure S8. Analysis of p(A) site mapping data.**

(A and B) Principal component analysis (PCA) plot of U937 Poly(A)-Click-sequencing data from samples of the control (0h) and cells treated with PMA for 6 hours (A) or 24 hours (B). Each dot represents a sample and each color represents the time of PMA treatment. (C and D) Volcano plots illustrating changes in gene expression counts and PAC counts. They were generated by plotting the $\log_2$ fold change values vs $p_{Adj}$ value for 6 hour (left panel) and 24 hour (right panel) PMA-treated U937 cells with respect to control undifferentiated ones for differential gene expression counts (C) and poly(A)-cluster counts (D). The gene counts for 6h and 24h collapse all mapped p(A) sites within one gene into one number and provides differential gene expression data, and PACs are defined as groupings of p(A) sites that are found within 25 nucleotides of one another. The gene counts in each PAC are used to determine relative changes in site selection between samples. Differential Gene or PAC count is defined as >1.5 fold change with $p_{Adj}$ <0.1 (independent hypothesis weighting).
